# Nutrient-sensitive histone propionylation regulates T cell fate and function

**DOI:** 10.64898/2026.10.01.756002

**Authors:** Alison Jaccard, Kelly Rome, Qihua Yang, Shuo Zhang, Noah Ford, Adam Chatoff, Tran Ngoc Van Le, Michael Noji, Andrea Andress Huacachino, Stephanie Stransky Lauar, Carlos Pondevida, Krittin Trihemasava, Laura V. Pinheiro, Sanat Bhadsalve, Eun Kyung Ko, Charles Salinas, Mariola M. Marcinkiewicz, Brian C. Capell, Golnaz Vahedi, Simone Sidoli, Roddy S. O’Connor, Will Bailis, Nathaniel W. Snyder, Kathryn E. Wellen

## Abstract

Tissue microenvironments differ widely in nutrient composition; yet how local nutrient conditions shape epigenetic programs that direct immune cell fate is poorly understood. We identify histone lysine propionylation (Kpr) as a chromatin mechanism linking nutrient availability to T cell fate decisions. Proteomic analysis reveals that Kpr is broadly reduced in exhausted T cells. H3K18pr is enriched at stemness-associated loci in effector T cells and depleted upon exhaustion. Isoleucine and propionate supply propionyl-CoA in T cells, and restricted nutrient availability within the tumor microenvironment limits histone Kpr, promoting T cell exhaustion. Disrupting isoleucine catabolism in T cells further suppresses Kpr, without altering histone acetylation, and impairs T cell function and antitumor immunity in a manner rescued by propionate. Dietary supplementation of isoleucine and propionate increases Kpr in tumor infiltrating lymphocytes, limits exhaustion, and improves tumor control. These findings establish a microenvironment-chromatin axis crucial for antitumor immunity and amenable to nutritional intervention.

## Introduction

CD8⁺ T cells orchestrate immune protection against infection and cancer by adopting distinct functional states that balance effector activity, persistence, and memory formation. However, under conditions of chronic antigen stimulation, T cells progressively differentiate into a dysfunctional state known as T cell exhaustion. Exhausted T (T_EX_) cells exhibit impaired cytokine production, sustained inhibitory receptor expression, and altered metabolic, transcriptional and epigenetic programs that hinder control of chronic infection and tumors [1–3]. A defining feature of the TME is its distinct metabolic landscape, characterized by nutrient depletion and accumulation of immunosuppressive metabolites [4–7]. T cells must therefore navigate different metabolic geographies as they traffic between lymph nodes, where progenitor populations principally reside, and tumors, where terminal exhaustion predominates. Whether the nutrient composition of these anatomical niches is actively translated into chromatin programs controlling T cell fate and function remains a fundamental open question. This question has high biologic importance and translational potential. T cells activated via their endogenous T cell receptor (TCR) are stimulated in the lymph node whereas T cells redirected to tumor antigens via chimeric antigen receptors (CARs) undergo constitutive activation in the TME.

T cell activation and differentiation are closely connected to metabolism, and progression toward exhaustion is accompanied by metabolic rewiring, including impaired mitochondrial function and altered nutrient utilization with increased reliance on glucose [6, 8, 9]. Beyond cellular energetics, certain metabolites derived from nutrient catabolism are used as substrates of chromatin-modifying enzymes and can thereby influence transcriptional programs [10–12]. Recent work has shown that acetyl-CoA availability links nutrient inputs to histone acetylation and T cell fate, demonstrating that metabolic signals can be integrated into chromatin programs [13]. Yet the repertoire of nutrient-sensitive chromatin modifications is incompletely defined, and whether distinct nutrient environments such as the TME and the tumor-draining lymph node (dLN) impose chromatin states that differentially regulate T cell fate and function across compartments is unknown.

Here, we address this question by identifying histone lysine propionylation (Kpr) as a chromatin modification that links local nutrient availability to T cell fate and function. Kpr arises from the three-carbon acyl group donor propionyl-CoA (pr-CoA), generated from several propiogenic nutrients, including the branched-chain amino acid (BCAA) isoleucine and the short-chain fatty acid (SCFA) propionate, both of which we find to be differentially abundant in the tumor versus the tumor dLN microenvironments. Although histone Kpr is a broadly distributed chromatin modification detected across diverse cell types, including cancer and immune cells [14–16], its biological functions and the metabolic pathways that regulate its deposition remain largely unknown. Using histone proteomics, we find that Kpr is broadly reduced during T cell exhaustion, with prominent loss at H3K18. Genome-wide mapping reveals that H3K18pr marks genes associated with T cell stemness and effector potential in effector CD8⁺ T cells and is depleted at these loci during exhaustion. Moreover, H3K18pr occupied genomic loci only partially shared with H3K18ac, revealing a regulatory layer not fully captured by acetylation. Genetic disruption of isoleucine catabolism reduced histone Kpr, promoted exhaustion and impaired T cell antitumor function selectively in the TME but not in the tumor dLN. Restoring propiogenic nutrient availability within the TME through metabolic supplementation restored T cell histone Kpr and antitumor activity, providing direct evidence that the local nutrient environment governs histone Kpr and T cell fate. These findings establish the principle that distinct tissue nutrient landscapes impose differential chromatin states in T cells and highlight this nutrient-to-chromatin axis as a tunable determinant of antitumor immunity.

## Results

### Histone propionylation is reduced during T cell exhaustion

To identify chromatin modifications associated with the regulation of T cell exhaustion, we used an in vitro model of acute and chronic stimulation and performed histone proteomics comparing effector T (T_E_) and T_EX_ cells (Fig. S1a). One of the most striking differences was an enrichment of histone propionylation (Kpr) in T_E_ versus T_EX_ cells across multiple lysine residues (Fig. 1a, b). Reduced histone Kpr in T_EX_ was confirmed at H3K18 and H4K16 by immunoblotting, as well as by flow cytometry at H3K18 (Fig. 1c, d). In line with prior work showing that H3K27ac is suppressed in T_EX_ cells, this mark was also suppressed in this model (Fig. 1c) [13].

**Figure 1.**
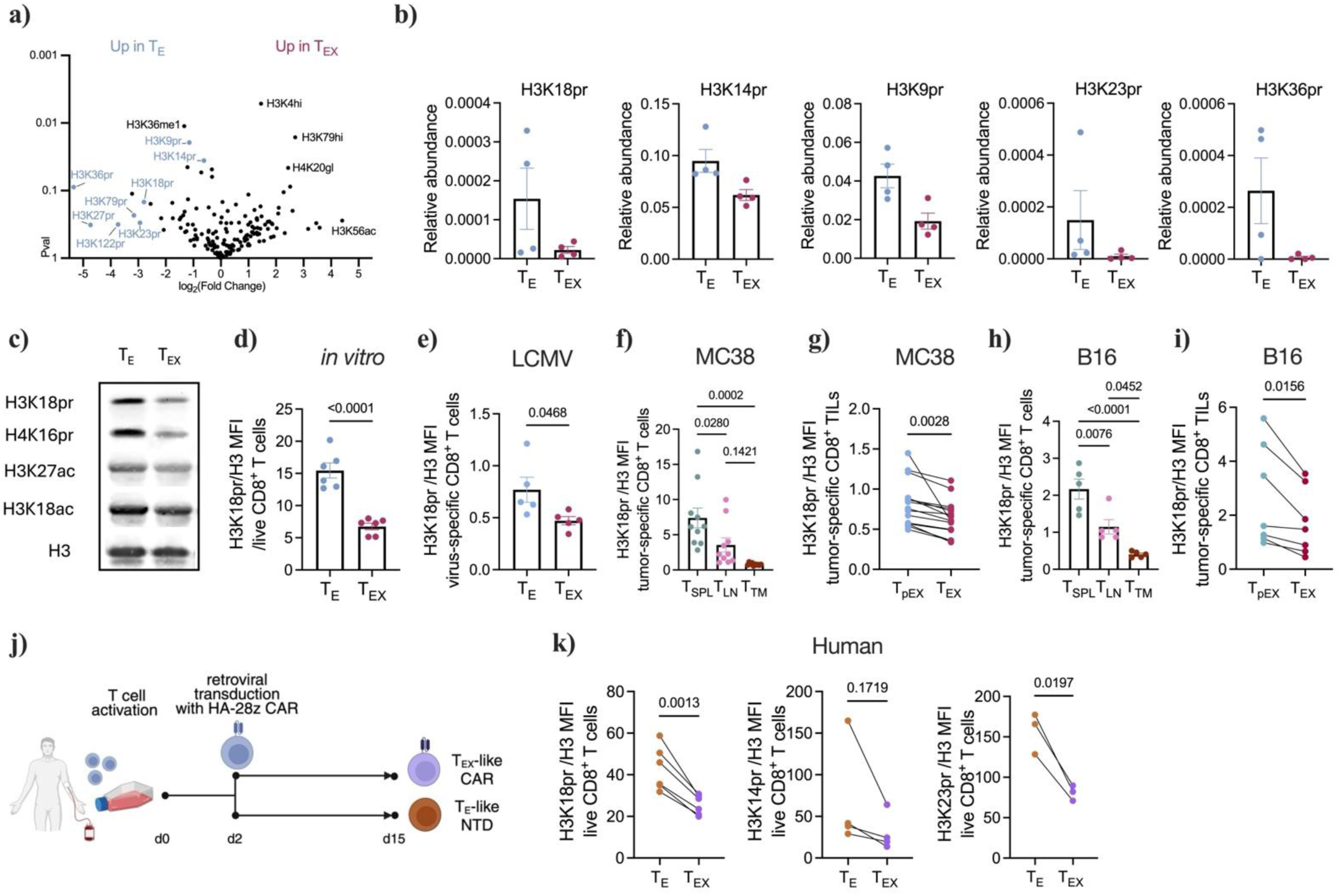
Histone propionylation is reduced during T cell exhaustion. **a)** Volcano plot of histone proteomics comparing effector (T_E_) and exhausted (T_EX_) T cells generated as described in Fig. S1a. **b)** Relative abundances of indicated histone modifications determined by histone proteomic analysis. **c)** Immunoblot of indicated histone marks in *in vitro*-generated T_E_ and T_EX_ cells. **d)** H3K18pr levels normalized to total H3 in *in vitro*-generated T_E_ and T_EX_ cells, measured by flow cytometry. **e)** H3K18pr levels normalized to total H3 in T_E_ (KLRG1^+^; CD127^-^) and T_EX_ (PD1^+^; TCF1^-^) P14 cells isolated from mice infected with LCMV Armstrong and Clone 13 respectively. **f)** H3K18pr levels normalized to total H3 in P14 cells isolated from the spleen (SPL), draining lymph node (LN) and tumor (TM) in MC38-gp33-bearing mice. **g)** H3K18pr levels normalized to total H3 in T_pEX_ (PD1^+^; TCF1^+^) and T_EX_ (PD1^+^; TCF1^-^) exhausted P14 TILs. **h)** H3K18pr levels normalized to total H3 in OT-1 cells isolated from the spleen (SPL), draining lymph node (LN) and tumor (TM) in B16-OVA-bearing mice. **i)** H3K18pr levels normalized to total H3 in T_pEX_ (PD1^+^; TCF1^+^) and T_EX_ (PD1^+^; TCF1^-^) exhausted OT-1 TILs. **j)** Schematic of a human CAR T cell exhaustion model driven by tonic signaling. Two days after activation, T cells were transduced with a GD2-specific CD28ζ CAR having a high affinity for GD2 (HA-28z) and maintained in culture for a total of 15 days. **k)** H3K18pr, H3K14pr and H3K23pr levels normalized to total H3 in non-transduced (NTD) T_E_ and HA-28z CAR T_EX_ cells, measured by flow cytometry. Data are shown as mean ± SEM. Statistical significance was determined using unpaired (d, e) or paired (g, i, k) Student’s t-test and by one-way analysis of variance (ANOVA) with Tukey’s multiple-comparison test (f, h).

To determine whether histone Kpr is also reduced in T_EX_ cells *in vivo*, we first analyzed antigen-specific P14 CD8^+^ T_E_ and T_EX_ cells isolated from mice infected with lymphocytic choriomeningitis virus (LCMV) Armstrong (Arm) and Clone 13 (Cl 13) respectively (Fig. S1b). We further focused on H3K18pr due to its consistent and robust reduction upon exhaustion and to specific antibody availability. T_EX_ cells generated from chronic viral infection (Cl 13) displayed lower histone H3K18pr than T_E_ cells arising during acute infection (Arm), as well as lower H3K27ac but not H3K18ac levels (Fig. 1e, Fig. S1c). Next, to test if T cell histone Kpr also varied in cancer settings, we used murine models of colon carcinoma (MC38-gp33) and melanoma (B16-OVA) with adoptive transfer of antigen-specific P14 and OT-1 T cells respectively (Fig. S1d, e). In both tumor models, H3K18pr levels were the lowest in antigen-specific tumor infiltrating CD8^+^ T cells (TILs), intermediate in T cells infiltrating the draining lymph node (dLN) and highest in splenic T cells (Fig. 1f, h). This gradient correlated with exhaustion states, as TILs exhibited the most terminally exhausted phenotype (Fig. S1f). Consistently, within tumors, terminally exhausted T (T_EX_) cells displayed lower histone H3K18pr than progenitor exhausted T (T_pEX_) cells (Fig. 1g, i). Finally, in a human model of CAR T cell exhaustion driven by tonic signaling (Fig. 1j, [17], we found that exhausted T cells expressing a HA-28z CAR had diminished histone Kpr compared to non-transduced (NTD) T cells, which lack tonic signaling and do not become exhausted (Fig. 1k, Fig. S1g). In this model, both H3K18ac and H3K27ac were also lower in the T_EX_ cells (Fig. S1h). Taken together, these results demonstrate that histone Kpr is reduced during T cell exhaustion across models, including in murine models of viral infection and of malignancies, as well as *in vitro* in both mouse and human T cells.

### Propionyl-CoA in T cells is supplied by isoleucine and propionate metabolism, which are altered in T_EX_ cells

Histone Kpr originates from the transfer of a propionyl group from pr-CoA to a lysine residue on a histone. Because previous work in other cell types demonstrated that intracellular pr-CoA pools positively correlate with histone Kpr levels [18, 19], we asked whether altered pr-CoA metabolism might explain the reduced histone Kpr observed in T_EX_ cells. Indeed, we measured a significant reduction in T_EX_ cell pr-CoA abundance (Fig. 2a), consistent with diminished histone Kpr levels, suggesting that altered T cell pr-CoA metabolism could be a mechanism for histone Kpr regulation during exhaustion.

**Figure 2.**
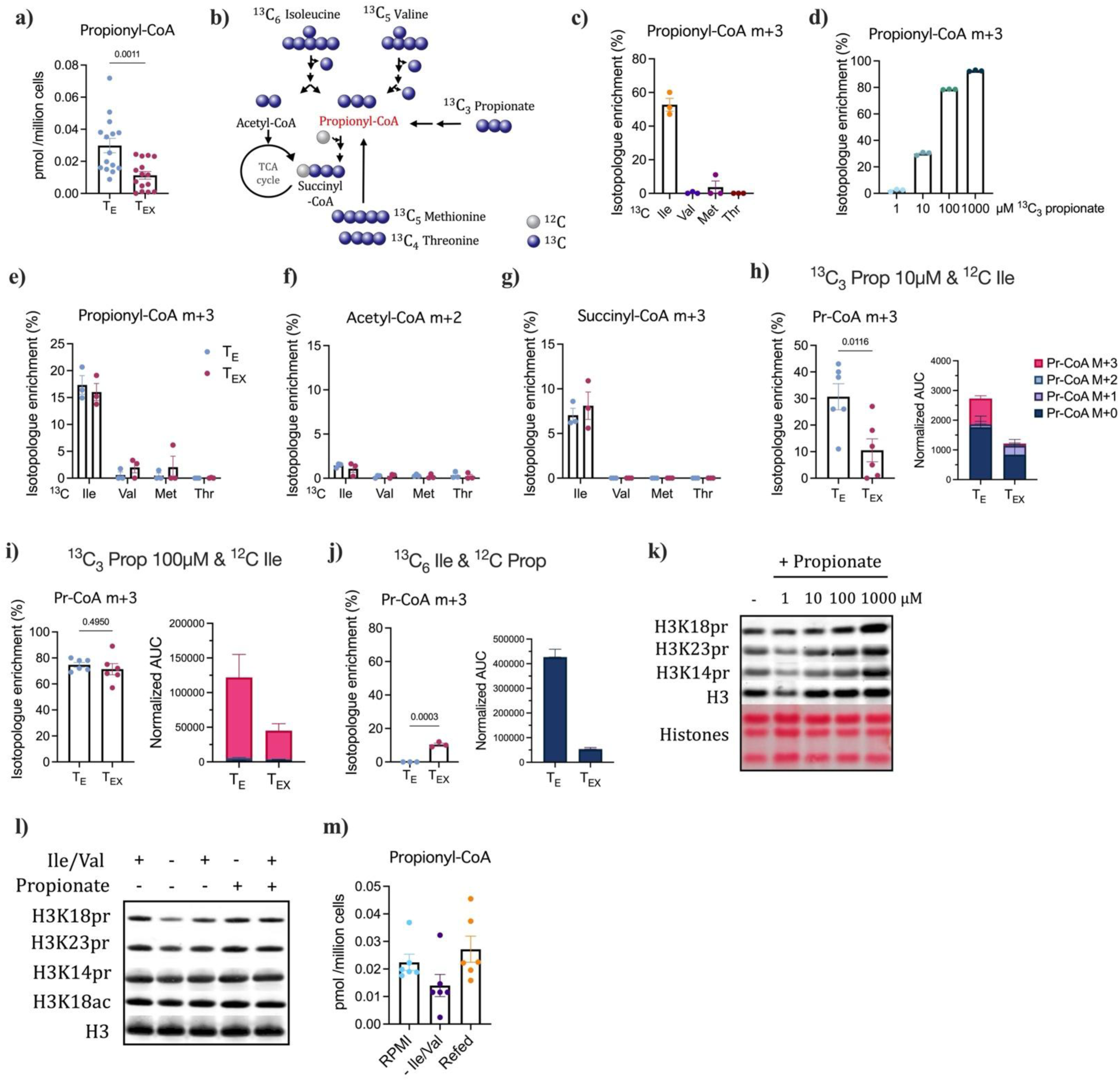
Histone Kpr is fueled by BCAA and propionate metabolism, which are altered upon T cell exhaustion. **a)** Pr-CoA abundance in picomoles per million cells in *in vitro*-generated T_E_ and T_EX_ cells measured by LC-MS. **b)** Schematic of carbon labeling contribution from indicated substrates to pr-CoA, acetyl-CoA and succinyl-CoA. **c)** Percent labeling of pr-CoA m+3 from 2h tracing with indicated substrates in CD8^+^ T cells at day 3 post activation. **d)** Percent labeling of pr-CoA m+3 from 2h tracing with [^13^C_3_]-propionate at indicated concentrations **e-g)** Percent labeling of pr-CoA m+3 (e), acetyl-CoA m+2 (f) and succinyl-CoA m+4 (g) from 2h tracing with indicated substrates in T_E_ and T_EX_ cells at day 8 post activation. **h and i)** Percent m+3 labeling of pr-CoA following 2h tracing with [^13^C_3_]-propionate at 10µM (h) or 100µM (i) in the presence of unlabeled isoleucine (582µM), and corresponding normalized area-under-the-curve quantification in T_E_ and T_EX_ cells. **j)** Percent m+3 labeling of pr-CoA following 2h tracing with [^13^C_6_]-isoleucine in the presence of unlabeled propionate (100µM), and corresponding normalized area-under-the-curve quantification in T_E_ and T_EX_ cells. **k)** Immunoblot of indicated histone marks in CD8^+^ T cells upon propionate supplementation at indicated concentrations. **l)** Immunoblot of indicated histone marks in CD8^+^ T cells upon Ile/Val deprivation and/or propionate supplementation. **m)** pr-CoA abundance in picomoles per million cells in CD8^+^ T cells upon indicated conditions measured by LC-MS. Data are shown as mean ± SEM; statistical significance was determined by paired Student’s t test (a, h-j).

Pr-CoA is a three-carbon acyl group donor that can be formed by the catabolism of several nutrient sources, including the BCAAs isoleucine and valine, the amino acids methionine and threonine and the short-chain fatty acid propionate [20]. To determine the contribution of these nutrients to pr-CoA generation in T cells, we used ^13^C-labelled substrates and measured their incorporation into pr-CoA (m+3 isotopologue) (Fig. 2b). Among potential amino acid sources of pr-CoA, activated CD8^+^ T cells primarily generate pr-CoA from Ile catabolism (Fig. 2c). Ile minimally labelled acetyl-CoA, succinyl-CoA and other TCA cycle intermediates, indicating that other carbon sources predominantly supply those intermediates (Fig. S2a, b). While propionate is not present in standard T cell culture conditions, when supplemented, propionate was incorporated into pr-CoA in a concentration-dependent manner, identifying this short-chain fatty acid as an additional pr-CoA source in T cells (Fig. 2d).

T_E_ and T_EX_ cells similarly relied on Ile as a primary pr-CoA source, with minimal percentage-wise contributions from other propiogenic substrates tested, and no significant difference in acetyl-CoA or succinyl-CoA labelling between the two populations (Fig 2e, f, g, Fig. S2c). However, T_EX_ cells showed impaired ability to use low concentrations of propionate (10µM) for pr-CoA synthesis (Fig. 2h). In contrast, at a higher propionate concentration (100µM), T_EX_ cells incorporated it into pr-CoA at similar percentages as T_E_ cells, despite their total pr-CoA pool remaining lower (Fig. 2i). Notably, at this higher propionate concentration, both T_E_ and T_EX_ cells preferentially utilized propionate over Ile to make pr-CoA (Fig. 2j). Consistent with a cell-intrinsic contribution to reduced pr-CoA in T_EX_ cells, analysis of published transcriptomic datasets from *in vitro* and *in vivo* exhaustion models revealed that *Acss1* and *Acss2*, two acyl-CoA synthetases with reported activity toward propionate, though with lower efficiency than for acetate [21], were downregulated in T_EX_ relative to T_E_ cells (Fig. S2e). *Acss3*, the primary pr-CoA synthetase in other tissues, was not detected in CD8^+^ T cells (Fig. S2e). These results suggest that reduced capacity for propionate utilization contributes to impaired pr-CoA synthesis in T_EX_ cells.

Since isoleucine and propionate serve as the main nutrient precursors for T cell pr-CoA generation, we tested whether their availability regulates histone Kpr. Indeed, propionate supplementation dose-dependently increased histone Kpr at several lysines (Fig. 2k). Conversely, Ile/Val deprivation diminished histone Kpr, which could be restored by propionate supplementation (Fig. 2l). The pr-CoA pool was impacted by these substrate manipulations in a pattern consistent with the histone Kpr changes (Fig. 2m). Notably, Ile/Val and propionate availability similarly influenced human CAR T cell histone Kpr (Fig. S2d).

Together, these findings uncover Ile-and propionate-derived pr-CoA as a metabolic node that bridges specific nutrient inputs to chromatin regulation via histone Kpr, and reveal that T_EX_ cells exhibit impaired propionate metabolism, pr-CoA synthesis and reduced intracellular pr-CoA abundance, which correlates with reduced histone Kpr.

### Nutrient availability controls histone propionylation and regulates T cell fate and function

To understand if this reduction in histone Kpr might regulate T cell fate decisions, we first mapped genome-wide chromatin localization of H3K18pr in T_E_ and T_EX_ cells generated as described in Fig. S1a using CUT&RUN. We detected an increased number of H3K18pr peaks unique to T_E_ cells and primarily located at promoters and distal intergenic regions (Fig. 3a, b), in line with higher Kpr levels observed in T_E_ and progenitor cells (Fig. 1). Gene ontology analysis of peaks present in T_E_ but lost in T_EX_ cells revealed H3K18pr enrichment at loci regulating T cell progenitor function, including Wnt signaling, notably *Tcf7*, *Lef1* and *Tcf4* (Fig. 3c-e) [22]. Although propionylation at K18 overlapped with acetylation at many loci in T_E_ cells, as would be expected for two marks associated with active chromatin, it also had distinct genomic deposition patterns at other loci (i.e., peaks were called for H3K18pr but not H3K18ac at certain loci). Gene ontology of H3K18pr-enriched loci identified Wnt signaling, mitophagy and Golgi transport pathways, supporting a model in which the regulation of propionylation is at least in part distinct from acetylation (Fig. 3e, f, Fig. S3a-c). As expected, the active chromatin mark H3K27ac was more abundant at progenitor genes including *Il2ra*, *Sell* and *Bach2* in T_E_ cells, as opposed to T_EX_ cells where this mark was increased at exhaustion-specific genes like *Pdcd1* and *Havcr2*, but not *Tox* (Fig. S3d, e). We also observed an increase in the repressive mark H3K27me3 at progenitor genes (*Tcf7*) in T_EX_ cells, whereas it was reduced at exhausted-associated loci (*Tox*) as previously reported [23] (Fig. S3e). These findings show that H3K18pr is remodeled during T cell exhaustion, with prominent losses of the modification at loci associated with a progenitor T cell phenotype.

**Figure 3.**
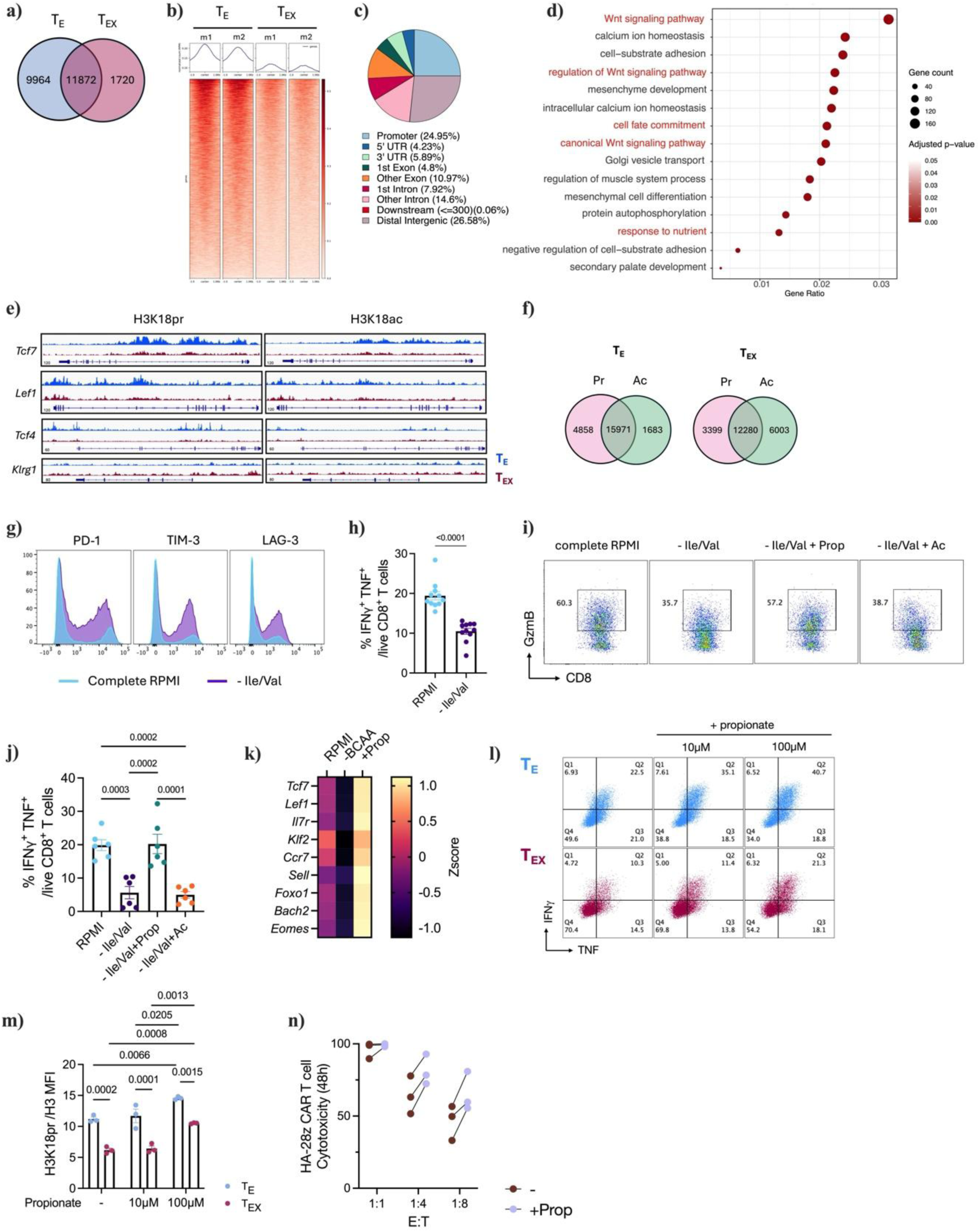
Nutrient conditions control histone propionylation and T cell fate and function. **a)** Venn diagram representing the overlap between H3K18pr peaks in T_E_ and T_EX_ cells. **b)** Heat map showing signal intensity of unique T_E_ H3K18pr peaks in two replicates of T_E_ and T_EX_ cells. **c)** Genomic distribution of H3K18pr peaks in T_E_ cells **d)** Gene ontology analysis on the H3K18pr peaks unique in T_E_ cells. **e)** Genome tracks at indicated loci showing H3K18pr and H3K18ac occupancy in T_E_ and T_EX_ cells. **f)** Venn diagrams representing the overlap between H3K18pr and H3K18ac peaks in T_E_ and T_EX_ cells. **g)** Representative histograms of PD-1, TIM-3 and LAG-3 expression in CD8^+^ T cells cultured in complete RPMI (fed) or under Ile/Val deprivation conditions. **h)** Percentage of IFNγ^+^ and TNF^+^ cells among fed or Ile/Val-deprived CD8^+^ T cells after restimulation with PMA/ionomycin. **i)** Representative dot plots of Granzyme B expression by fed and Ile/Val-deprived CD8^+^ T cells with propionate or acetate supplementation following restimulation. **j)** Percentage of IFNγ^+^ and TNF^+^ cells among fed or Ile/Val starved CD8^+^ T cells with propionate or acetate supplementation following restimulation. **k)** Heatmap showing expression of selected differentially expressed genes (FDR<0.05) in fed, starved and propionate-supplemented starved CD8^+^ T cells. **l)** Representative dot plot of IFNγ^+^ and TNF^+^ T_E_ and T_EX_ cells following 6-day supplementation with 10 or 100µM propionate and PMA/ionomycin restimulation. **m)** H3K18pr levels normalized to total H3 in *in vitro*-generated T_E_ and T_EX_ cells following 6-day supplementation with 10 and 100µM propionate, measured by flow cytometry. **n)** Cytotoxicity of HA-28z CAR T cells at indicated effector-to-target (E:T) ratios measured after a 48 h co-culture, with control NaCl (-) or sodium propionate (+Prop) supplementation throughout the duration of the experiment. Data are shown as mean ± SEM; statistical significance was determined by unpaired Student’s t test (h) and by one-way (j) or two-way ANOVA with Tukey’s multiple-comparison test (l, m).

To test whether pr-CoA synthesis plays a regulatory role in T cell differentiation and functionality, we examined the impact of nutrient perturbations that alter pr-CoA abundance and Kpr. Ile/Val starvation promoted inhibitory receptor expression and impaired effector cytokine production, hallmarks of T cell exhaustion (Fig. 3g, h, Fig. S3f, g). Remarkably, propionate supplementation restored granzyme B and cytokine production, whereas acetate, a SCFA that is not a precursor to pr-CoA, failed to rescue this defect, supporting a role for pr-CoA in regulating CD8^+^ T cell function (Fig. 3i, j). Furthermore, transcriptomic analyses showed downregulation of stemness-associated genes upon BCAA starvation, that was similarly restored by propionate (Fig. 3k). Thus, pr-CoA availability couples nutrient conditions to a stemness-like chromatin state and functional T cell outputs.

We next asked whether restoring pr-CoA in T_EX_ cells could reverse low histone Kpr and functional defects. Supplementing T_E_ cells with propionate for six days (Fig. S3h) dose-dependently boosted pr-CoA abundance and effector cytokine production (Fig. 3l, Fig. S3h-j) and was accompanied by an increase in H3K18pr as well (Fig. 3m). Strikingly, in T_EX_ cells, long-term propionate supplementation (100 μM) (Fig. S3h) increased cellular pr-CoA abundance, and restored cytokine production and histone Kpr levels (Fig. 3l, m, Fig. S3i, j). Furthermore, in the human model of CAR T cell exhaustion (Fig. 1j), propionate supplementation during *in vitro* expansion enhanced HA-28z CAR T cell cytotoxicity along with cellular pr-CoA abundance (Fig. 3n, Fig. S3k).

Together, these data identify a key role for Ile and propionate nutrient availability in promoting histone H3K18pr deposition at progenitor-specific genes, thereby reprogramming the effector versus exhausted T cell epigenetic state and T cell function.

### BCAA and propionate metabolism regulate T cell Kpr, fate and function

Next, to specifically test whether Ile catabolism regulates histone Kpr and T cell function, we disrupted the pathway using CRISPR/Cas9-mediated deletion of ACAT1, the enzyme in Ile catabolism that directly generates pr-CoA (Fig. 4a, Fig. S4a). ACAT1-deficient CD8^+^ T cells displayed reduced pr-CoA but not acetyl-CoA abundance, and reduced ^13^C_6_-Ile incorporation into pr-CoA (Fig. 4b-d). Consistent with reduced pr-CoA abundance, ACAT1 deletion reduced H3K18pr but not H3K18ac and impaired effector cytokine production, both of which were restored by propionate supplementation (Fig. 4e-g, Fig. S4b, c). These findings were recapitulated in both CRISPR-deleted BCKDHA and CD8^+^ T cells isolated from mice with a T cell-specific deletion of DBT (Dbt^fl/fl^ ; CD4-cre mice), confirming that, similar to Ile starvation, genetic disruption of the BCAA catabolic pathway impairs pr-CoA metabolism, histone Kpr and T cell functionality (Fig. S4d-j).

We hypothesized that ACAT1-deficient T cells would exhibit impaired ability to control tumor growth. To test the role of Ile catabolism in antitumor immunity, we adoptively transferred P14 ACAT1-deficient (KO) or Rosa-targeted control (WT) CD8^+^ T cells into MC38-gp33 tumor-bearing mice (Fig. 4h). Transfer of WT P14 cells conferred antitumor effects compared to untreated mice (Fig. S4k), whereas ACAT1-deficient P14 cells had significantly reduced capacity to control tumor growth following adoptive cell transfer (ACT) (Fig. 4i, j). ACAT1-deficient P14 cells exhibited reduced H3K18pr, in line with a model in which loss of Ile-dependent histone Kpr is associated with terminal exhaustion and impaired antitumor T cell function (Fig.4k). Consistently, animals receiving ACAT1-deficient P14 cells had reduced intratumoral T cell accumulation (Fig. 4l). Among the transferred TILs, the ACAT1-deficient P14 cell phenotype was shifted toward a more terminally exhausted and less progenitor exhausted phenotype, even though total numbers of both populations were decreased in the tumor (Fig. 4m-o, Fig. S4l). ACAT1-deficient cells also displayed impaired IFNγ and TNF cytokine production upon *ex vivo* restimulation (Fig.4p). Intriguingly, in contrast to that in the tumors, the tumor dLN contained comparable numbers of ACAT1-deficient and WT P14 cells (Fig.4q), with no differences in the number of progenitor and terminally exhausted populations, cytokine production, or histone Kpr levels (Fig.4r-u).

**Figure 4.**
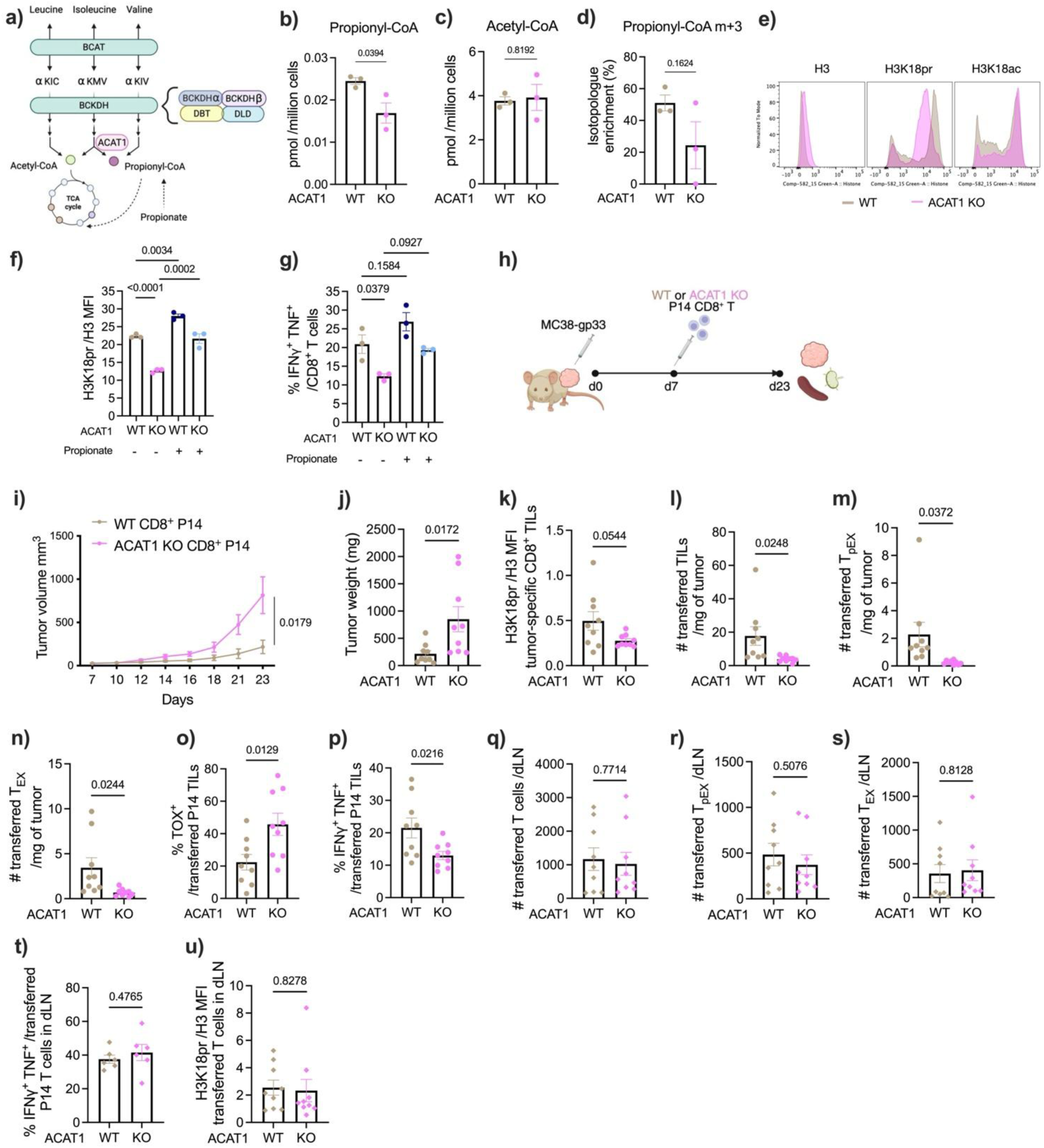
BCAA and propionate metabolism regulate T cell Kpr, fate and function. **a)** Schematic of BCAA and propionate metabolic pathways contributing to pr-CoA generation. **b and c)** pr-CoA (b) and acetyl-CoA (c) abundances (pmol per million cells) in Rosa-targeted control (WT) and ACAT1-deficient (KO) CD8^+^ T cells **d)** Percent m+3 labeling of pr-CoA following 2h tracing with [^13^C_6_]-isoleucine in Rosa control (WT) and ACAT1-deficient (KO) CD8^+^ T cells. **e)** Representative histograms of H3, H3K18pr and H3K18ac staining in Rosa control (WT) and ACAT1-deficient (KO) CD8^+^ T cells, measured by flow cytometry. **f)** H3K18pr levels normalized to total H3 in Rosa control (WT) and ACAT1-deficient (KO) CD8^+^ T cells with or without 100µM propionate supplementation. **g)** Percentage of IFNγ^+^ and TNF^+^ cells among Rosa control (WT) and ACAT1-deficient (KO) CD8^+^ T cells, with or without 100µM propionate supplementation following restimulation. **h)** Rosa-targeted control (WT) or ACAT1-deficient (KO) P14 CD8^+^ T cells were adoptively transferred into B6 mice 7 days after MC38-gp33 tumor implantation. **i and j)** Tumor growth curve (i) and tumor weights (j) of mice bearing MC38-gp33 tumors and transferred with Rosa control (WT) or ACAT1-deficient (KO) P14 CD8^+^ T cells. **k)** H3K18pr levels normalized to total H3 in Rosa control (WT) and ACAT1-deficient (KO) P14 CD8^+^ TILs **l-n)** Numbers of total P14 CD8^+^ TILs (l), progenitor exhausted (PD1^+^ TCF1^+^ T_pEX_) (m) and terminally exhausted (PD1^+^ TCF1^-^ T_EX_) (n) per milligram of tumor in Rosa control (WT) and ACAT1-deficient (KO) mice. **o)** Percentage of TOX^+^ cells among Rosa control (WT) and ACAT1-deficient (KO) P14 TILs **p)** Percentage of IFNγ^+^ and TNF^+^ cells among Rosa control (WT) and ACAT1-deficient (KO) P14 CD8^+^ TILs after *ex vivo* restimulation. **q-s)** Numbers of total (q), T_pEX_ (PD1^+^ TCF1^+^) (r) and T_EX_ (PD1^+^ TCF1^-^) (s) P14 CD8^+^ T cells in the tumor draining lymph node (dLN) of mice receiving Rosa control (WT) and ACAT1-deficient (KO) cells. **t)** Percentage of IFNγ^+^ and TNF^+^ cells among Rosa control (WT) and ACAT1-deficient (KO) P14 CD8^+^ T cells isolated from the dLN following *ex vivo* restimulation. **u)** H3K18pr levels normalized to total H3 in Rosa control (WT) and ACAT1-deficient (KO) P14 CD8^+^ T cells isolated from the dLN. Data are shown as mean ± SEM; statistical significance was determined by unpaired Student’s t test (b-d, i, j-u) and by one-way ANOVA with Tukey’s multiple-comparison test (f, g).

Therefore, disruption of Ile catabolism to pr-CoA promoted T cell exhaustion in the TME, correlating with loss of histone Kpr and tumor control. These data identify Ile catabolism as a critical metabolic pathway sustaining histone Kpr via pr-CoA and supporting the CD8^+^ T cell antitumor response.

### Nutrient composition in microenvironments dictates T cell fate and function via histone propionylation

Because ACAT1-deficient T cells exhibited a phenotype consistent with increased terminal exhaustion and reduced histone Kpr in tumors, but not in the dLN, we hypothesized that histone Kpr might depend on the local nutrient environment. This model predicts that availability of pr-CoA precursors differs across microenvironments such as the TME and the tumor dLN. To test this hypothesis, we profiled metabolite composition in plasma, tumor-and dLN-interstitial fluids (TIF and LIF). As previously reported, glutamine and arginine were depleted in TIF compared to plasma, lactate was elevated in TIF (Fig.5a-c, [4, 8]), and glucose was highest in LIF but had similar concentrations in plasma and TIF (Fig.5d). Notably, both propionate and Ile were significantly lower in abundance in TIF compared to LIF, though also more abundant than in plasma (Fig.5e, f). Intriguingly, the propionate concentration was approximately 100 μM in LIF, a concentration sufficient to sustain T_EX_ cell pr-CoA pools, histone Kpr and cytokine production in the absence of Ile (Fig.2). The propionate concentration in TIF, on the other hand, was in a lower concentration range where T_EX_ cells may be unable to sustain pr-CoA synthesis, histone Kpr and effector function (Fig.2). These findings suggest that the tumor dLN constitutes a metabolic niche where Ile and propionate availability allows for sustained histone Kpr in T cells, whereas lower abundance of these pr-CoA precursors in the TME might promote loss of Kpr and terminal exhaustion.

To directly test this possibility and the role of Ile-and propionate-dependent histone Kpr in antitumor immunity, we supplemented tumor-bearing mice with sodium propionate in the drinking water at time of ACT with ACAT1-deficient or control P14 cells (Fig.5g). While propionate supplementation did not affect body weight or propionate concentrations in plasma or LIF, it significantly elevated propionate abundance within TIF (Fig.5h, i, Fig. S5a, b). As observed in figure 4, ACAT1-deleted P14 T cells exhibited reduced ability to control tumor growth compared to WT P14 T cells (Fig. 5j, k, Fig. S5c). However, consistent with elevated TIF propionate levels, propionate supplementation increased TIL abundance and restored tumor control mediated by ACAT1-deficient P14 cells (Fig.5j-l). Notably, improved tumor growth control was also observed in mice receiving WT T cell ACT with propionate supplementation (Fig. 5j, k). Propionate supplementation also partially reversed exhaustion marker expression (i.e., TOX), restored IFNγ and TNF production, and boosted H3K18pr levels in both WT and ACAT1-deficient TILs, while it had no effect on antigen-specific T cells infiltrating the tumor dLN and spleen (Fig.5m-o, Fig. S5d-h). These data indicate that loss of Ile-dependent Kpr and T cell functionality upon ACAT1 deletion can be rescued by sufficient propionate supplementation, which also restores histone Kpr independently of Ile catabolism, limits terminal exhaustion, and re-establishes T cell antitumor activity.

**Figure 5.**
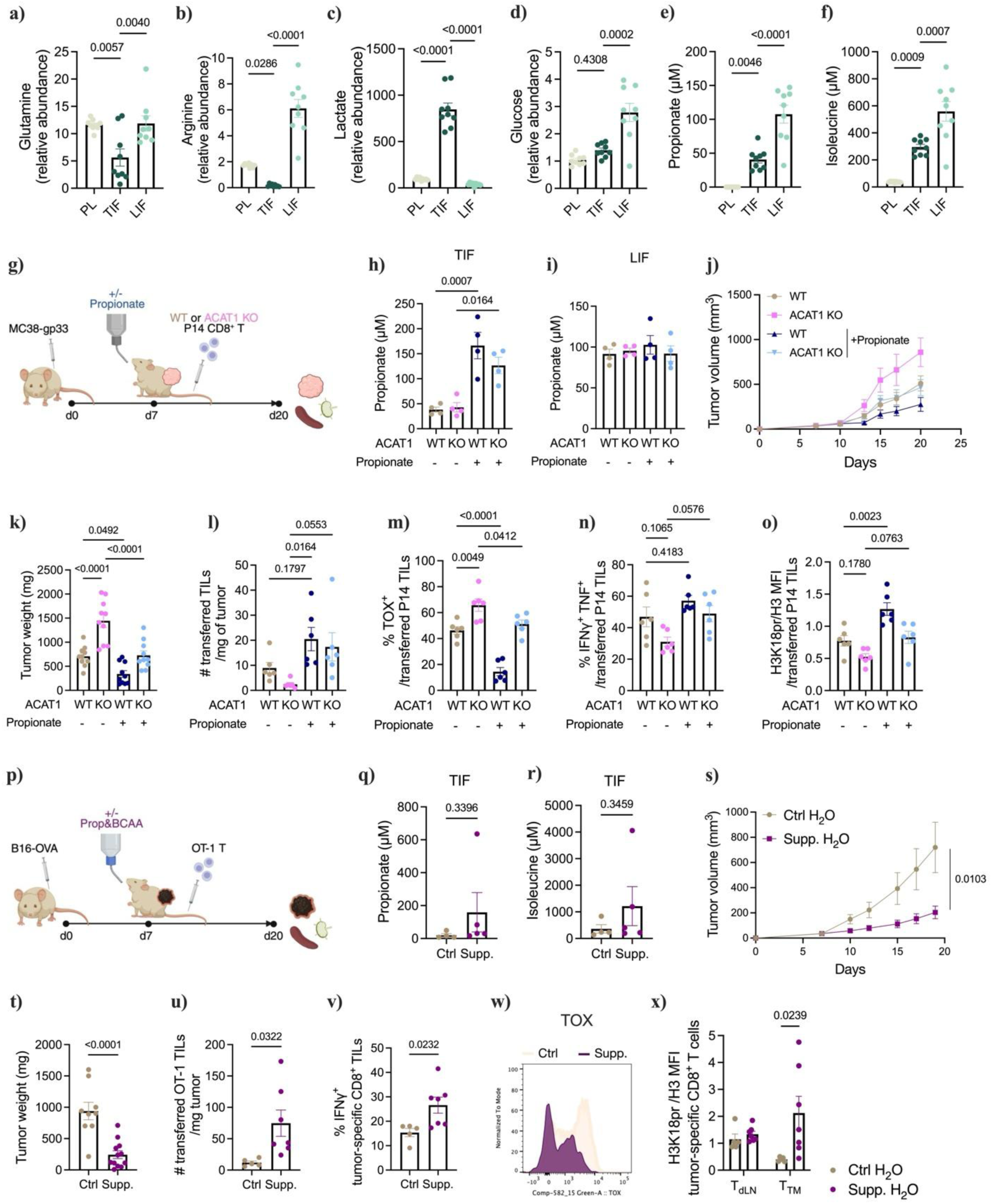
Nutrient composition in microenvironments dictates T cell fate and function via histone propionylation. **a-d)** Relative abundances of glutamine (a), arginine (b), lactate (c) and glucose (d) in plasma (PL), tumor-interstitial fluid (TIF) and lymph node-interstitial fluid (LIF) of MC38-gp33 tumor-bearing mice measured by LC-MS. **e-f)** Concentrations of propionate (e) and isoleucine (f) in plasma (PL), tumor-interstitial fluid (TIF) and lymph node-interstitial fluid (LIF) of MC38-gp33 tumor-bearing mice measured by LC-MS. **g)** Rosa control (WT) or ACAT1-deficient (KO) P14 CD8^+^ T cells were adoptively transferred into B6 mice 7 days after MC38-gp33 tumor implantation and sodium propionate (150mM) or control NaCl was administered in the drinking water beginning at the time of adoptive cell transfer. **h and i)** Propionate concentrations in TIF (h) and LIF (i) from control or propionate-supplemented tumor-bearing mice receiving Rosa control (WT) or ACAT1-deficient (KO) P14 CD8^+^ T cells. **j and k)** Tumor growth curve (j) and tumor weights (k) of mice bearing MC38-gp33 tumors and receiving control or propionate-supplemented drinking water. **l)** Numbers of P14 CD8^+^ TILs per mg of tumor in indicated groups. **m)** Percentage of TOX^+^ cells among P14 CD8^+^ TILs of indicated groups **n)** Percentage of IFNγ^+^ and TNF^+^ cells among P14 CD8^+^ TILs of indicated groups upon *ex vivo* restimulation. **o)** H3K18pr levels normalized to total H3 in P14 CD8^+^ TILs of indicated groups. **p)** Rosa control (WT) or ACAT1-deficient (KO) OT-1 cells were adoptively transferred into B6 mice 7 days after B16-OVA tumor implantation and propionate (150 mM) and Ile/Val (0.8% Ile and 1.12% Val) were supplemented in the drinking water beginning at the time of adoptive cell transfer. **q and r)** Concentrations of propionate (q) and isoleucine (r) in TIF of B16-OVA tumor-bearing mice measured by LC-MS. **s and t)** Tumor growth curve (s) and tumor weights (t) of mice bearing B16-OVA tumors and transferred with OT-1 T cells, receiving control or propionate/BCAA-supplemented drinking water. **u)** Numbers of OT-1 TILs per mg of tumor in control and supplemented mice. **v)** Percentage of IFNγ^+^ cells among OT-1 TILs of control and supplemented mice upon *ex vivo* restimulation. **w)** Representative flow cytometry histogram of TOX expression in control and supplemented OT-1 TILs. **x)** H3K18pr levels normalized to total H3 in control and supplemented OT-1 T cells isolated from the dLN and tumor. Data are shown as mean ± SEM; statistical significance was determined by unpaired Student’s t test (q-v) and by one-way (a-f, h, i, k-o) or two-way (x) ANOVA with Tukey’s multiple-comparison test.

We also noted that raising propionate availability in TIF also enhanced the antitumor activity of WT CD8^+^ T cells, as it further reduced tumor growth, weight and TOX expression (Fig.5j, k, m), while increasing TIL numbers, effector function and histone Kpr (Fig.5I, n, o). To further evaluate whether boosting pr-CoA precursor availability in the TME can preserve histone Kpr in TILs and enhance antitumor immunity in a second model, we supplemented B16-OVA melanoma tumor-bearing mice with propionate and Ile/Val in the drinking water, along with OT-1 ACT (Fig.5p). We confirmed that nutrient supplementation in the drinking water raised propionate and Ile TIF abundances in these settings (Fig.5q, r), although there was some variability. Plasma concentrations of Ile and Val, but not propionate, also increased, and body weight was unaltered (Fig. S5i-k). Propionate and Ile/Val supplementation triggered robust antitumor immunity, reducing tumor growth and end point tumor weight (Fig.5s, t). Nutrient supplementation also significantly boosted TIL numbers and TIL IFNγ production upon *ex vivo* restimulation, and reduced TIL TOX expression (Fig.5u, v, w, Fig. S5l-p). Finally, increasing propionate and Ile availability in the TME restored histone Kpr in TILs to levels comparable to those in tumor dLN-infiltrating T cells (Fig.5x). Consistent with our previous findings, nutrient supplementation did not impact the OT-1 cells infiltrating the tumor dLN (Fig.5x, Fig. S5q, r).

Together, these results establish that Ile and propionate metabolism sustain histone Kpr to promote T cell antitumor function and limit T_EX_ differentiation. Loss of nutrient-dependent histone Kpr promoted T cell dysfunction, whereas nutrient supplementation preserved histone Kpr, facilitating resilience against exhaustion and antitumor immunity. These findings define a nutrient-metabolite-chromatin axis in which environmental nutrient availability regulates pr-CoA metabolism, histone Kpr and ultimately T cell fate and antitumor function.

## Discussion

While the metabolic hostility of the TME is well recognized as a barrier to T cell function, whether specific nutrient limitations are actively encoded into chromatin, rather than solely impairing T cell metabolism, has remained unclear. Our findings provide a mechanistic framework: the TME is relatively depleted of propiogenic nutrients compared to the tumor-dLN, and this nutrient landscape is translated into differential histone Kpr that regulates T cell fate across compartments.

We found that the BCAA isoleucine and the SCFA propionate sustain histone Kpr in T cells via the intermediate pr-CoA, and that their limited availability in the TME leads to histone Kpr loss at several stemness-associated loci, contributing to T cell terminal exhaustion and loss of antitumor capacity. Genetic disruption of Ile catabolism reduces T cell histone Kpr and accelerates terminal exhaustion specifically in the TME but not in the tumor dLN, where propionate availability remains sufficient, and restoring propionate availability to TILs rescues both Kpr and antitumor function. These data provide evidence that the nutrient environment controls histone Kpr and regulates T cell fate.

These findings fit within broader evidence that metabolic pathways regulate T cell differentiation by controlling the availability of metabolites that support chromatin modification [24–33]. While prior studies focused largely on acetyl-CoA–dependent histone acetylation and S-adenosylmethionine-dependent histone methylation, our findings identify histone Kpr as a novel nutrient-responsive regulator of T cell state governed by distinct metabolic enzymes and nutrient sources from those regulating histone acetylation. Specifically, we found that Kpr is particularly sensitive to propiogenic nutrients, which are present at limiting levels for the regulation of Kpr within the TME, contributing to T cell exhaustion and limiting antitumor immunity.

Although histone Kpr has similar biochemical properties to histone lysine acetylation [18], and several histone acetyltransferases can utilize both acetyl-CoA and propionyl-CoA as acyl donors [19], our findings suggest that these two chromatin marks exhibit distinct mechanisms of regulation. Indeed, we observed that H3K18pr occupies specific genomic loci, only partially overlapping with H3K18ac. Nutrient-dependent regulation of pr-CoA availability selectively alters H3K18pr without impacting H3K18ac, indicating that specificity may come from distinct upstream regulation by metabolic state and nutrient availability. These findings imply that even though chromatin acylations might share enzymatic machineries, they can be distinctly regulated by differential metabolite availability. Notably, we found that T cell effector function depends on histone Kpr sustained by propiogenic substrate availability, which is limited in the TME compared to lymph nodes. Unlike acetyl-CoA, which can be generated via multiple metabolic pathways, pr-CoA is derived from more restricted propiogenic sources, making it particularly sensitive to nutrient abundance. Consequently, fluctuations in nutrient availability may selectively influence cellular pr-CoA pools and modulate histone propionylation independently of histone acetylation. Histone Kpr may thus function as a chromatin mark that encodes specific metabolic inputs rather than simply mirroring acetylation patterns. While the Ile catabolism pathway generating pr-CoA is well established, the enzyme responsible for propionate activation in T cells remains to be identified, as ACSS3, the primary propionyl-CoA synthetase in other tissues [34], is not expressed in CD8^+^ T cells. Other acyl-CoA synthetase family members, including ACSS1 and ACSS2, may fulfill this role, though this requires direct investigation.

Consistent with the idea that histone Kpr is regulated at the level of metabolite availability, our findings show that tissue nutrient landscapes can shape Kpr levels in T cells. We observed that lymph nodes constitute metabolic niches enriched in the propiogenic substrates Ile and propionate. Notably, lymph nodes are the primary sites where stem-like and progenitor T cells reside, populations that displayed the highest levels of histone Kpr. These findings suggest that the nutrient-rich environment of lymphoid tissues supports elevated cellular pr-CoA pools, thereby sustaining Kpr deposition and progenitor potential and T cell function. An open question raised by our findings is why lymph nodes are enriched in propiogenic nutrients. The mechanisms that establish these nutrient landscapes remain unclear and elucidating the sources of these metabolites in tumors and in lymphoid tissues and how their availability is regulated will be an important direction for future work.

Finally, our findings raise the possibility that boosting histone Kpr through augmenting pr-CoA synthesis could represent a nutritional strategy to promote functional T cell states for immune therapies. In support of this, we show that supplementation with propionate, a metabolite that reaches the TME, increases cellular pr-CoA availability, is generally well tolerated and even used as a food-safe additive, may represent a tractable approach with translational potential. Previous reports showed that SCFA supplementation can boost CD8^+^ T cell effector cytokine production. These studies focused on other SCFAs (e.g. pentanoate and butyrate), and similarly found that propionate could boost cytokine production, though they did not explore downstream mechanisms [35, 36]. Similarly, increased BCAA availability has recently been linked to improved T cell antitumor immunity, and to synergize with immune checkpoint blockade therapy in murine cancer models, even though the mechanistic understanding is also limited [37, 38].

Our finding that propionate supplementation enhances CAR T cell cytotoxicity *in vitro* suggests that boosting pr-CoA availability may represent a tractable strategy to improve CAR T cell therapy. We speculate that CAR T cells may be particularly susceptible to propiogenic nutrient depletion in the TME, as unlike conventional T cells which can egress into the tumor dLN where propiogenic nutrients are more abundant, CAR T cells undergo constitutive activation within the TME and may have limited opportunity to replenish pr-CoA pools through exposure to nutrient-replete lymphoid niches. Directly testing whether propionate supplementation enhances CAR T cell persistence and antitumor function *in vivo* represents an important and clinically relevant future direction.

Although manipulation of nutrient and microbiota-derived metabolite availability, including BCAAs and SCFAs, has been reported to modulate T cell function through metabolic reprogramming [35, 37, 38] and acetyl-CoA–dependent epigenetic programs [13, 39], our findings identify histone propionylation as a distinct and context-dependent chromatin mechanism that directly bridges local metabolite availability and T cell fate decisions. Our study thus identifies a key mechanism by which propionate and Ile/Val enhance T cell effector function via increasing histone Kpr. More broadly, these results highlight the potential of targeting nutrient–epigenetic interactions to enhance immune responses and improve cancer immunotherapy.

Together, these findings establish histone propionylation as a nutrient-responsive chromatin mechanism linking tissue metabolic environments to T cell states, and representing a tunable axis for shaping T cell immunity.

## Acknowledgements

We thank all members of the Wellen and the Snyder laboratories for helpful discussions and critical feedback. We thank Mathias Wenes for his valuable input and for reviewing the manuscript. We thank Michael Michieletto for kindly sharing some reagents and reviewing the manuscript. We would like to thank Irfan Asangani and Erick Mitchell for use of NextSeq for CUT&RUN sequencing, and Simone Park and John Wherry for providing OT-1 mice. Flow cytometry data for this manuscript were generated in the Penn Cytomics and Cell Sorting Shared Resource Laboratory at the University of Pennsylvania (RRID:SCR_022376). Penn Cytomics is partially supported by the Abramson Cancer Center NCI Grant (P30 CA016520). Figures were created in part using Biorender.com. This work was funded by grants from the Ludwig Institute for Cancer Research and a Mark Endeavor Award to K.E.W. and NCI R01-CA-292937 to K.E.W. and N.W.S., and A.J. was supported by a Swiss National Science Foundation (SNSF) Postdoc.Mobility fellowship. K.R was supported by NIH T32CA009140, M.C.N. was supported by F31CA261041 and L.V.P. was supported by an AFCRI RISE postdoctoral fellowship.

## Author contributions

A.J., K.E.W. and N.W.S. conceptualized the project, designed experiments, interpreted and discussed results, and wrote the manuscript. K.E.W. and N.W.S. acquired funding. A.J., K.R., Q.Y., N.F., A.C., T.N.V.L., M.N., A.A.H., S.S.L., C.P., K.T., L.V.P., S.B., E.K.K., C.S. and M.M. performed experiments. S.Z. discussed and analyzed sequencing results. S.S.L. and S.S. analyzed histone proteomic results. K.E.W., N.W.S, W.B., R.S.O, S.S., G.V., B.C. supervised work. All authors reviewed and approved the manuscript.

## Material & Methods

### Mice & cell lines

C57BL/6 (B6) (CD45.2) mice were purchased from The Jackson Laboratory. P14 and Dbt conditional knockout (cKO) mice were obtained from the Bailis laboratory, and OT-I mice were obtained from the Wherry laboratory. All animal experimentation was conducted in accordance with protocols approved by the University of Pennsylvania (UPenn) Institutional Animal Care and Use Committee (IACUC). Mice were maintained under specific pathogen-free conditions on a 12 h light/ 12 h dark cycle at 21-23°C, and fed standard rodent chow (LabDiet, 5010) and water, unless otherwise indicated. Male and female mice between 6 and 10 weeks of age were used for experiments and were age-and sex-matched for adoptive transfer experiments. MC38-gp33 and B16-OVA melanoma cells were obtained from the Bailis laboratory and were cultured in DMEM, containing 10% fetal bovine serum (FBS), 1% Penicillin/Streptomycin (P/S, Gibco 15070-063), and 1% L-glutamine (Q, Gibco 25030-081). Cell lines were routinely tested for mycoplasma contamination by Polymerase Chain Reaction (PCR) and confirmed to be mycoplasma-free.

### Mouse and human CD8^+^ T cell isolation

Single-cell suspensions were prepared from spleens and peripheral lymph nodes of C57BL/6 mice by mechanical dissociation through 70-μm cell strainers, followed by red blood cell lysis (Qiagen). Mouse CD8⁺ T cells were isolated using the EasySep™ Mouse CD8⁺ T Cell Isolation Kit (STEMCELL Technologies). Human CD8⁺ T cells were isolated using the Human CD8⁺ T Cell Isolation Kit and LS magnetic columns (Miltenyi Biotec) according to the manufacturer’s instructions.

### Mouse T cell culture

Isolated CD8⁺ T cells were activated on plate-bound anti-CD3/CD28 antibodies (BioLegend) at 1 × 10⁶ cells mL⁻f in RPMI 1640 (Gibco) supplemented with 10% fetal bovine serum (FBS), 10 mM HEPES, 1 mM sodium pyruvate, 1× non-essential amino acids, 2 mM L-glutamine, 50 μM β-mercaptoethanol, 1% penicillin-streptomycin, and recombinant human IL-2 (100 IU mL⁻f; PeproTech). For generation of *in vitro* effector (T_E_) and exhausted (T_EX_) CD8⁺ T cells, activated cells were either cultured without further stimulation until day 8 to generate T_E_ cells or restimulated with anti-CD3/CD28 every other day until day 8 to generate T_EX_ cells.

### CRISPR-Cas9 RNP Nucleofection of Primary CD8⁺ T Cells

CRISPR-Cas9 ribonucleoprotein (RNP) complexes were generated by combining target-specific crRNAs directed against Acat1 or the Rosa26 safe-harbor locus (Integrated DNA Technologies) with tracrRNA and recombinant Cas9 protein. crRNA duplexes were formed by incubation at 95°C for 5 min followed by cooling to room temperature, after which Cas9 protein (QB3 Berkeley MacroLab) was added and incubated for 15 min prior to nucleofection. Activated P14 CD8⁺ T cells were washed with PBS, resuspended in P3 Nucleofector Solution (Lonza), mixed with Cas9-RNP complexes, and nucleofected using a 4D-Nucleofector (Lonza) with program DN100. Following nucleofection, cells were recovered in antibiotic-free complete medium and cultured for downstream experiments. ACAT1 deletion efficiency was confirmed by immunoblot analysis of ACAT1 protein expression.

crRNA target sequences:

Mouse Rosa26: ACTCCAGTCTTTCTAGAAGA

Mouse Acat1: GUCCCAUACGUAAUGAGCAGGUUUUAGAGCUAUGCU

Mouse Bckdha: TACGGCAACGTGAATGACCC

### Retrovirus Production

10 million 293GP cells were plated in 20mL of DMEM supplemented with 10% FBS, 10 mM HEPES and 1× penicillin– streptomycin–glutamate (Gibco) onto Poly-D Lysine-coated 15cm plates and cultured at 37°C overnight. Transfection was performed by mixing a room-temperature solution of 1.5 ml Opti-MEM (Gibco) + 120 μl Lipofectamine 3000 (Invitrogen) (solution 1) with a second solution of 1.5 ml Opti-MEM + 60uL P3000 + 10 μg RD114 packaging plasmid DNA + 20 μg MSGV retroviral plasmid of interest (solution 2) by slow dropwise addition of solution 2 to solution 1. The combined solution 1 and 2 mixture was incubated for 30 min at room temperature, after which the medium was fully replaced on 293GP cells, then 3 ml of the combined solution was added to the plates in a slow, dropwise manner before being returned to culture at 37°C. The next day, the culture medium was replaced. Forty-eight hours after transfection, the viral supernatant was collected from the cells and the culture medium was replaced. Supernatant collection was repeated at 72 hours. After each supernatant collection, the supernatant was spun down (3000 RPMx 10 minutes) to remove cells and debris, and frozen at −80 °C for future use.

### Retroviral Transduction

For HA.28z CAR T experiments, T cells were transduced on days two and three post-activation. Briefly, 24-well non tissue culture-treated plates were coated with 500uL of 20ug/mL retronectin (Takara) overnight at 4C. The next day, plates were washed with PBS and plates were blocked with 3% BSA in PBS for 10 min. Retroviral supernatant was added and the plate was centrifuged at 32°C for 2 hours and 20 min. Viral supernatants were then removed and 2.5 e5 T cells were added to each well at a concentration of 5e5 cells/mL in T cell medium.

### HA.28z cell culture

T cells were thawed in warm water after removal from liquid nitrogen and then washed with T cell medium (AIM-V (Gibco) supplemented with 5% FBS, 10 mM HEPES, 1× penicillin–streptomycin–glutamate and 100 U ml^−1^ recombinant human IL-2 (Peprotech). Primary human T cells from three donors were activated using CD3/28 dynabeads (Gibco) at a 2:1 bead:T cell ratio at 1e6 cells/mL. T cells were transduced with retrovirus two and three days after activation per previously established protocols. Four days after initial activation, beads were removed and cells were resuspended in 2mL total of fresh AIMV media per retroviral transduction well. Cells were counted and fed to 500,000 cells/mL on day 7 post-activation and every other day thereafter until day 15.

### LCMV infection model

C57BL/6 mice were infected with either Lymphocytic Choriomeningitis Virus (LCMV) Armstrong (2 × 10⁵ PFU, intraperitoneal) or LCMV Clone 13 (4 × 10⁶ PFU, retro-orbital). On the day of infection, P14 CD8⁺ T cells were isolated and adoptively transferred into recipient mice. Flow cytometry analyses were performed on blood and spleen samples 8 days following LCMV Armstrong infection or 28 days following LCMV Clone 13 infection.

### Tumor models and adoptive cell transfer

Recipient C57BL/6 mice were subcutaneously engrafted with 5 × 10⁵ gp33-expressing MC38 or OVA-expressing B16 tumor cells. Seven days after tumor implantation, mice were randomized based on tumor volume and 1 × 10⁶ P14 or OT-I CD8⁺ T cells were adoptively transferred intravenously. Tumor growth was monitored every 2 days using digital calipers, and tumor volumes were calculated according to the formula V = π(d² × D)/6, where d represents the minor tumor axis and D the major tumor axis. At the indicated time points, tumors were harvested and dissociated into single-cell suspensions using collagenase and DNase (Sigma-Aldrich). Tumors, tumor draining lymph nodes and spleens were mechanically dissociated through 70 μm cell strainers. Single-cell suspensions were stained with fluorochrome-conjugated antibodies and analyzed by flow cytometry.

### Nutrient supplementation

At the time of adoptive cell transfer, mice were randomized to receive drinking water supplemented with either sodium propionate (150 mM), or sodium propionate (150 mM) supplemented with isoleucine and valine (0.8% and 1.12%, respectively), or a sodium-matched NaCl control. Supplemented water was provided ad libitum throughout the duration of the experiment and refreshed weekly. Mouse body weights were monitored weekly to assess tolerability. Metabolite concentrations in plasma and indicated tissues were quantified by LC-MS to confirm systemic and tissue-specific nutrient exposure.

### Flow Cytometry

Fluorochrome-conjugated antibodies were all from Biolegend or Thermo Fisher Scientific and non-conjugated antibodies were from Cell Signaling. Staining for flow cytometry was done in PBS with 2% FBS and 2mM EDTA, at 4°C in the dark. The Life/Dead Fixable Blue Cell Stain Kit (Thermo Fisher Scientific) was used to mark dead cells. The Foxp3 staining kit (Thermo Fisher Scientific) was used for intranuclear staining, and BD Bioscience kit was used for intracellular cytokine staining, following the manufacturer’s instructions. Acquisition was done on LSR II, Fortessa and Symphony flow cytometers with FACSDiva Software.

### Western Blotting

Whole cell lysis was done in RIPA lysis buffer (50 mM TrisHCl pH8, 150 mM NaCl, 1% Triton X 100, 0.5% Sodium deoxycholate, 0.1% SDS) to which 500µM of fresh protease and phosphatase inhibitors (Phosstop, Sigma) were added. Histones were isolated by centrifugation at 6,500 g for 10 min at 4°C and the pH was neutralized with 2 M NaOH at 1/10 of the volume of the supernatant. Proteins were quantified with BCA protein assay kit (Thermo Fisher Scientific, 2309). Equal amounts of protein extracts or of histone extracts were denatured 5 minutes at 95 C in 4X laemmli sample buffer (Bio-Rad, 161-0747) containing β-mercaptoethanol, followed by separation on gradient SDS-PAGE gels (Invitrogen, NP0322BOX) and transferred to nitrocellulose membranes (Bio-Rad, 1620115). Membranes were stained with Ponceau S Staining solution (Cell Signaling, 59803S) and blocked with 5% BSA in PBST 1 hour at RT. After transfer, blocking of unspecific binding sites was done in 5% BSA in PBST and incubation with primary antibodies (Cell Signaling, PTM BIO, Abcam, Milipore, CST, Proteintech, Origene) was done overnight at 4°C. The next day, membranes were incubated with secondary fluorescently labelled anti-rabbit or anti-mouse antibodies (LicorBio) and developed with a LI-COR Odyssey Clx system.

### Cytotoxicity Assays

Twenty thousand NALM6 cells expressing GFP, GD2, and CBG luciferase were cocultured with D15 HA.28Z CAR T cells at the indicated E:T ratios in 200 µL R10 in flat-bottomed 96-well assay plates for 48 hours. Each donor and condition was plated in triplicate. D-luciferin was added to each coculture well in 10 µL to reach a final concentration of 150 µg/mL and incubated at 37°C for 10 min in the dark. Luminescence was read on a BioTek Synergy HTX plate reader at 37°C. Cytotoxicity was calculated as (1-(LumCoculture/LumControl))*100. Values were plotted as the average of triplicate wells.

### Interstitial Fluid Isolation

Tumor interstitial fluid (TIF) was isolated as previously described with minor modifications [4]. Briefly, tumors were rapidly excised, weighed, rinsed in isotonic saline, blotted dry, and placed onto 70 μm nylon cell strainers positioned within 50 mL conical tubes. Samples were centrifuged at 300 × g for 10 min at 4°C, and the interstitial fluid recovered from the bottom of the collection tube was immediately frozen in liquid nitrogen and stored at −80°C until analysis. The same procedure was used for the isolation of interstitial fluid from tumor-draining lymph nodes. Plasma was collected at the time of sacrifice and processed in parallel for metabolomic analyses.

### Stable isotope tracing

For stable isotope tracing experiments, activated CD8⁺ T cells were cultured in custom RPMI tracing media prepared from RPMI powder (United States Biological) and supplemented with 10% dialyzed fetal bovine serum (dFBS). Tracing media were supplemented with glucose, glutamine, and all amino acids except the nutrient being traced. Cells were washed with PBS and incubated in tracing medium containing the indicated isotopically labelled nutrient for 2 h at 37°C and 5% CO₂. For BCAA tracing experiments, RPMI lacking isoleucine or valine was supplemented with [U-f³C]-isoleucine or [U-f³C]-valine. For methionine and threonine tracing experiments, custom RPMI lacking methionine or threonine, respectively, was supplemented with [U-f³C]-methionine or [U-f³C]-threonine. For propionate tracing experiments, [U-f³C]-propionate was added to the tracing medium at the indicated concentration. Unlabeled control samples were cultured under identical conditions using the corresponding unlabeled nutrients. Following incubation, cells were rapidly washed with ice-cold saline and metabolites were extracted for LC-MS analysis. Isotopologue distributions and fractional enrichments were corrected for natural isotope abundance using FluxFix [41].

### Acyl-Coenzyme A measurement

For extraction of acyl-CoAs, CD8^+^ T cells were isolated from mice spleens and lymph nodes and activated as described above. At day 3, cells were collected, counted and cultured for 2h with isotopically labelled nutrients (Cambridge Isotope Laboratories) in isoleucine-free RPMI containing 10% dialysed FBS. Acyl-CoA quantification and isotope tracing were measured by liquid chromatography-high resolution mass spectrometry as previously described in detail [42, 43]. Briefly, cell pellets were resuspended in 1mL 10% (w/v) trichloroacetic acid in water for extraction. Samples were homogenized by using a probe tip sonicator in 0.5 second pulses 30 times then centrifuged at 17,000 x g for 10 min at 4°C. Supernatant was purified by solid phase extraction (SPE) cartridges (Oasis HLB 10 mg, Waters) that were conditioned with 1 mL of methanol and 1 mL of water. Acid-extracted supernatants were loaded onto the cartridges and washed with 1 mL of water. Acyl-CoAs were eluted with 1 mL of 25 mM ammonium acetate in methanol and evaporated to dryness under nitrogen. Samples were resuspended in 50 µL of 5% (w/v) 5-Sulfosalicyilic acid and were analyzed by a Vanquish Duo UHPLC System (Thermo Fisher Scientific) coupled to a Thermo QExactive Plus instrument with a HESI II probe operating in positive ion mode. For enrichment analysis, isotopically labeled isoleucine enrichment into acyl-CoAs was determined using FluxFix [41].

### Amino acid and TCA cycle metabolite quantification

For polar metabolomics of BCAA and tricarboxylic acid (TCA) cycle metabolites, samples were quenched in 1 ml pre-chilled at −80 °C 80:20 methanol:water (vol/vol) and spiked with 50 µl isotopically labelled amino acids (10 µM) (Cambridge Isotope Laboratories). After vortexing for 1 min, samples were returned to −80 °C for 1 hour, centrifuged at 17,000g for 10 min at 4 °C, and the supernatant was transferred to a deep-well 96-well plate and evaporated to dryness under nitrogen gas. Samples were reconstituted in 50 µl, and 2 µl of the sample was injected from a 4°C autosampler into a ZIC-pHILIC 150 × 2.1 mm 5 µm particle size column (EMD Millipore) with a ZIC-pHILIC 20 × 2.1 guard column in a Vanquish Duo UHPLC system (Thermo Fisher Scientific) at 25 °C. Chromatography conditions were as follows: solvent A was acetonitrile; solvent B was 20 mM ammonium carbonate, 0.1% (vol/vol) ammonium hydroxide in water without pH adjustment, with a gradient of 0.5 min at 20% A, then a linear gradient from 20% to 80% B; 20–20.5 min: from 80% to 20% B; 20.5–28 min: hold at 20% B at a 0.150 mL/min−1 flow rate. Column elute was introduced to a Q Exactive Plus with a HESI II probe operating in polarity switching mode with full scans from 70 to 1,000 m/z with an insource fragmentation energy of 1.

### Propionate derivatization and quantification

Propionate was quantified by LC-MS following derivatization with 2-hydrazinoquinoline (HQ). Samples were extracted in ice-cold 80:20 methanol using the same method as Amino acid and TCA cycle metabolite quantification, and 10 μM [U-f³C₃]-propionate was used as an internal standard. Following clarification by centrifugation, 10 μL of extract was subjected to derivatization using 1 mM HQ in the presence of 1 mM triphenylphosphine (TPP) and 1 mM 2,2-dipyridyl disulfide (DPDS) in acetonitrile, with a total reaction volume of 110 μL. Samples were incubated at 60°C for 60 min, and the reaction was quenched with 80 μL optima grade water. Derivatized samples were clarified by centrifugation and only the supernatant was taken for LC-MS analysis. A Vanquish UPLC system (Thermo Fisher Scientific) coupled to a Exploris 480 mass spectrometer with a H-ESI source (Thermo Fisher Scientific) was used for propionate quantification. 1uL of the supernatant was injected onto a reverse-phase chromatography column, Acuity UPLC BEH C18 column, 1.7 μM, 2.1 x 50 mm (Waters). Chromatography conditions were as follows: Solvent A was 5 mM Ammonium Acetate in water, Solvent B was 5 mM ammonium acetate in 99.5% acetonitrile, 0.5% water, with starting conditions of 1% B at a flow rate of 0.200 mL/min and held for one minute. Then solvent B increased to 100% over 4 minutes at a flow rate of 0.250 mL/min. 100% B was maintained for 2 minutes at a flow rate of 0.300 mL/min. Solvent B was then reduced to 1% over 0.1 minutes at a flow rate of 0.250 mL/min. 1% B was maintained for one minute. The mass spectrometer operating conditions were 3,500 V spray voltage, 350 °C ion transfer tube temperature, 400 °C vaporizer temperature, and positive ion polarity. Both full scan detection from 50 to 500 m/z and data independent acquisition (DIA) detection with a center mass of 216.1131 m/z and 219.1232 m/z were used for quantification. Absolute propionate concentrations were determined using a standard curve generated from serial dilutions of unlabeled sodium propionate processed in parallel with experimental samples. Quantification was performed relative to the [U-f³C₃]-propionate internal standard to account for extraction efficiency, derivatization efficiency, and instrument variability.

### Histone proteomics: Histone extraction and derivatization and LC-MS/MS Acquisition and Analysis

Histones were extracted from cell pellets by incubation in 0.2 M sulfuric acid at 4°C followed by precipitation with trichloroacetic acid. Histone pellets were washed with acetone containing 0.1% HCl and subsequently with 100% acetone before air drying. Purified histones were resuspended in 100 mM ammonium bicarbonate (pH 8.0) containing 15–20% acetonitrile and subjected to two rounds of derivatization with propionic anhydride. Samples were then digested with trypsin in 50 mM ammonium bicarbonate (pH 8.0). Following digestion, peptide N-termini were derivatized by two additional rounds of propionylation with d10-propionic anhydride, which allows differentiation of endogenous propionylation. Samples were dried by vacuum centrifugation and stored at −80°C until analysis.

Prior to mass spectrometry analysis, samples were desalted using a 96-well plate filter (Orochem) packed with 1 mg of Oasis HLB C-18 resin (Waters). Briefly, samples were resuspended in 100 µl of 0.1% TFA and loaded onto the HLB resin, which was previously equilibrated with 100 µl of the same buffer. Following a wash with 100 µl of 0.1% TFA, peptides were eluted with 70 µl of a solution containing 60% acetonitrile and 0.1% TFA and then dried in a vacuum centrifuge.

Samples were resuspended in 10 µl of 0.1% TFA and analyzed using a Dionex RSLC Ultimate 300 (Thermo Scientific) coupled online to an Orbitrap Fusion Lumos (Thermo Scientific). Chromatographic separation was performed with a two-column setup consisting of a C-18 trap cartridge (300 µm ID, 5 mm length) and a picofrit analytical column (75 µm ID, 25 cm length) packed in-house with Repro-Sil Pur C18-AQ 3 µm reversed-phase resin. Histone peptides were separated using a 30 min gradient from 1% to 30% buffer B (buffer A: 0.1% formic acid, buffer B: 80% acetonitrile + 0.1% formic acid) at a flow rate of 300 nl/min.

The mass spectrometer was operated in data-independent acquisition (DIA) mode using isolation windows. Full MS scans were acquired over a mass range of 300-1100 m/z in the orbitrap at a resolution of 120,000 (at 200 m/z), with an AGC target of 5 x10^5^. MS/MS spectra were acquired in the orbitrap using sequential isolation windows of 50 m/z with an AGC target of 2 x10^5^ and an HCD collision energy of 30.

Histone peptides raw files were imported into EpiProfile 2.0 software [44] for data analysis. Peptide abundance was estimated from the area under the extracted ion chromatogram peaks. To calculate the relative abundance of post-translational modifications (PTMs), the summed signal intensity of all modified forms of a given histone peptide was defined as 100%, and the area corresponding to each individual modified form was normalized to this total. For isobaric peptide species, relative abundances were determined by averaging the ratios obtained from fragment ions that differed in mass between the two forms. The peptide tables generated by EpiProfile were exported to Microsoft Excel for further processing and downstream analyses.

### CUT&RUN

CUT&RUN was performed on effector and exhausted CD8⁺ T cells using the CUTANA CUT&RUN v5 Kit (EpiCypher) according to the manufacturer’s instructions with minor modifications optimized for primary T cells. Briefly, 5 × 10^5^ cells per sample were harvested and subjected to nuclei isolation. Nuclei were immobilized on activated concanavalin A-coated magnetic beads and incubated overnight at 4°C with antibodies against H3K18pr, H3K18ac, H3K27ac, H3K27me3, or rabbit IgG as a negative control. Two independent biological replicates were analyzed for each condition. Following antibody binding, nuclei were incubated with protein A/G–micrococcal nuclease (pAG-MNase; EpiCypher). Targeted chromatin digestion was initiated by addition of CaCl₂ and allowed to proceed for 2 h at 4°C. Digestion was terminated using Stop Buffer, and released chromatin fragments were recovered from the supernatant following magnetic separation. CUT&RUN DNA was purified using Monarch PCR & DNA Cleanup columns (New England Biolabs) and quantified using a Qubit dsDNA High Sensitivity assay. Sequencing libraries were generated using the NEBNext Ultra II DNA Library Prep Kit for Illumina (New England Biolabs). Purified DNA underwent end repair and A-tailing, followed by adaptor ligation and USER enzyme treatment. Libraries were amplified by PCR using indexed Illumina primers and purified using SPRIselect beads. Prior to sequencing on a NextSeq 2000 (Illumina), final library quality and fragment size distribution were assessed using an Agilent Bioanalyzer 2100 with a High Sensitivity DNA Chip (Agilent, 5067-4626).

### CUT&RUN analysis

Raw reads were aligned to mm39 reference genome using bowtie2 (v2.1.0) [45] with parameters: -q --local --very-sensitive --no-mixed -p 10 --no-unal --phred33. Alignments with mapping quality score less than 20 were filtered out using samtools (v1.9) [46]. Duplicates and blacklist regions [47] were removed using picard (https://github.com/broadinstitute/picard) and bedtools (v2.30.0) [48], respectively. Normalized tracks were generated using deepTools (v3.5.4) [49] bamCoverage with parameters: --normalizeUsing CPM. Peaks were called in each replicate over corresponding IgG control using MACS2 [50] with parameters: -f BAMPE -g mm -q 0.05 --broad --broad-cutoff 0.05 --max-gap 1000. We merged peaks from two replicates of each histone modification in T_E_ and T_EX_ cells, except that only replicate 1 (m1) was used for TEX H3K18ac as relative few peaks were called from replicate 2 (m2). Peak overlapping analysis was performed using bedtools. Peaks were annotated to nearest genes with “TxDb.Mmusculus.UCSC.mm39.knownGene” database using ChIPseeker package [51] and Gene Ontology (GO) was performed using clusterProfiler package [52].

### RNA sequencing analysis

Total RNA was isolated with the RNeasy Plus Mini Kit (Qiagen, 74134), according to the manufacturer’s protocol. RNA quality was confirmed using NanoDrop 2000 and RNA was sent to Novogene for standard non-directional library preparation and bulk mRNA-sequencing using Illumina NovaSeq X Plus, PE150 at a depth of 6Gb (20M PE reads) sequencing per sample.

Raw reads were pre-processed using fastp (v0.23.4) [53] with parameters: --detect_adapter_for_pe --cut_front --cut_tail. Preprocessed reads were aligned to mm10 reference genome with UCSC refGene annotation using STAR (v2.7.1a) [54] with parameters: --outFilterMismatchNmax 3 --outFilterMultimapNmax 1. Then, gene-level read counts were summarized using featureCounts (subread v2.0.6) [55]. For differential gene expression analysis, unwanted variation was first removed using RUVg of RUVseq package (v1.40.0) [56] with 2,000 least significant control genes called from DESeq2 (v1.46.0) [57] likelihood ratio test. Then, differentially expressed genes were called using DESeq2 with ashr (v2.2-63) [58] for log2 fold change shrinkage.

## Statistical analysis

A two-tailed unpaired Student’s t test was used to compare means between two independent groups, and comparisons involving three or more groups were analyzed using one-way ANOVA followed by Tukey’s post hoc test, unless indicated otherwise in the figure legends. All in vitro results shown reflect at least two independent experiments. Flow cytometry files (.fcs) were processed using FlowJo v10.10.0. Data are presented as mean ± SEM unless otherwise specified. GraphPad Prism version 10.6.0 (796) was used for statistical analyses. Sample sizes were not predetermined using formal power calculations, but are consistent with sizes used in related published articles. n indicates the number of biologically independent samples analyzed.

## Data availability

Histone proteomics data were deposited in the PRIDE (Pro-teomeXchange) database under accession: PXD079597. CUT&RUN and RNA-seq generated by this study have been deposited to Gene Expression Omnibus with accession number GSE335742 and GSE335739, respectively.

---

**Fig. S1.**
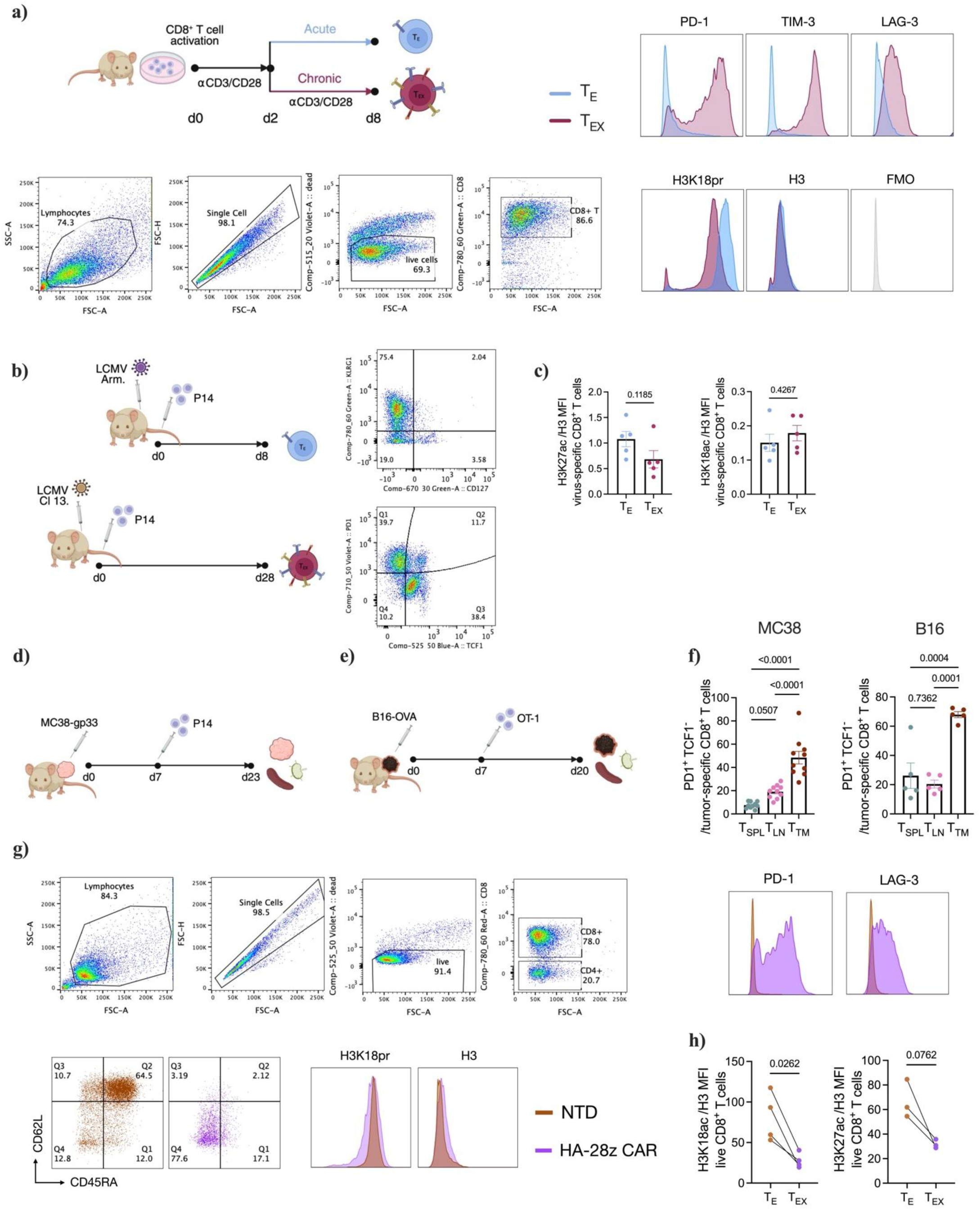
Histone propionylation is reduced during T cell exhaustion. a) CD8^+^ T cells were isolated and activated with αCD3/CD28 and IL-2. Two days post-activation, cells were either cultured without αCD3/CD28 to generate T_E_ cells, or restimulated with αCD3/CD28 every other day until day 8 to generate T_EX_ cells. Representative flow cytometry gating strategy and histograms of PD-1, TIM-3 and LAG-3 expression at day 8 in *in vitro*-generated T_E_ and T_EX_ cells. Representative flow cytometry histograms of H3K18pr and H3 levels in *in vitro*-generated T_E_ and T_EX_ cells, with FMO shown as negative control. **b)** B6 mice were adoptively transferred with P14 CD8^+^ T cells and infected with either LCMV Arm to generate T_E_ cells (KLRG1^+^; CD127^-^) or Cl13 to generate T_EX_ cells (PD1^+^; TCF1^-^). Representative flow cytometry plots showing gating of T_E_ and T_EX_ P14 cells. **c)** H3K27ac and H3K18ac levels normalized to total H3 in T_E_ (KLRG1^+^; CD127^-^) and T_EX_ (PD1^+^; TCF1^-^) P14 cells isolated from mice infected with LCMV Arm and Cl13 respectively. **d)** P14 CD8^+^ T cells were adoptively transferred into B6 mice 7 days after MC38-gp33 tumor implantation and tumor, tumor dLN and spleen were harvested at day 23. **e)** OT-1 T cells were adoptively transferred into B6 mice 7 days after B16-OVA tumor implantation and tumor, tumor dLN and spleen were collected at day 20. **f)** Percentage of PD1^+^; TCF1^-^ P14 cells isolated from the spleen (SPL), tumor draining lymph node (dLN) and tumor (TM) in MC38-gp33 (left) or B16-OVA (right) tumor-bearing mice. **g)** Representative flow cytometry gating strategy, plots and histograms of PD-1, LAG-3, CD62L and CD45RA expression and H3K18pr and H3 levels in NTD (T_E_) and HA-28z CAR (T_EX_) CD8^+^ T cells at day 15 post-activation. h) H3K18ac and H3K27ac levels normalized to total H3 in NTD T_E_ and HA-28z CAR T_EX_ cells, measured by flow cytometry. Data are shown as mean ± SEM. Statistical significance was determined using unpaired (c) or paired (h) Student’s t-test and by one-way ANOVA with Tukey’s multiple-comparison test (f).

**Fig. S2.**
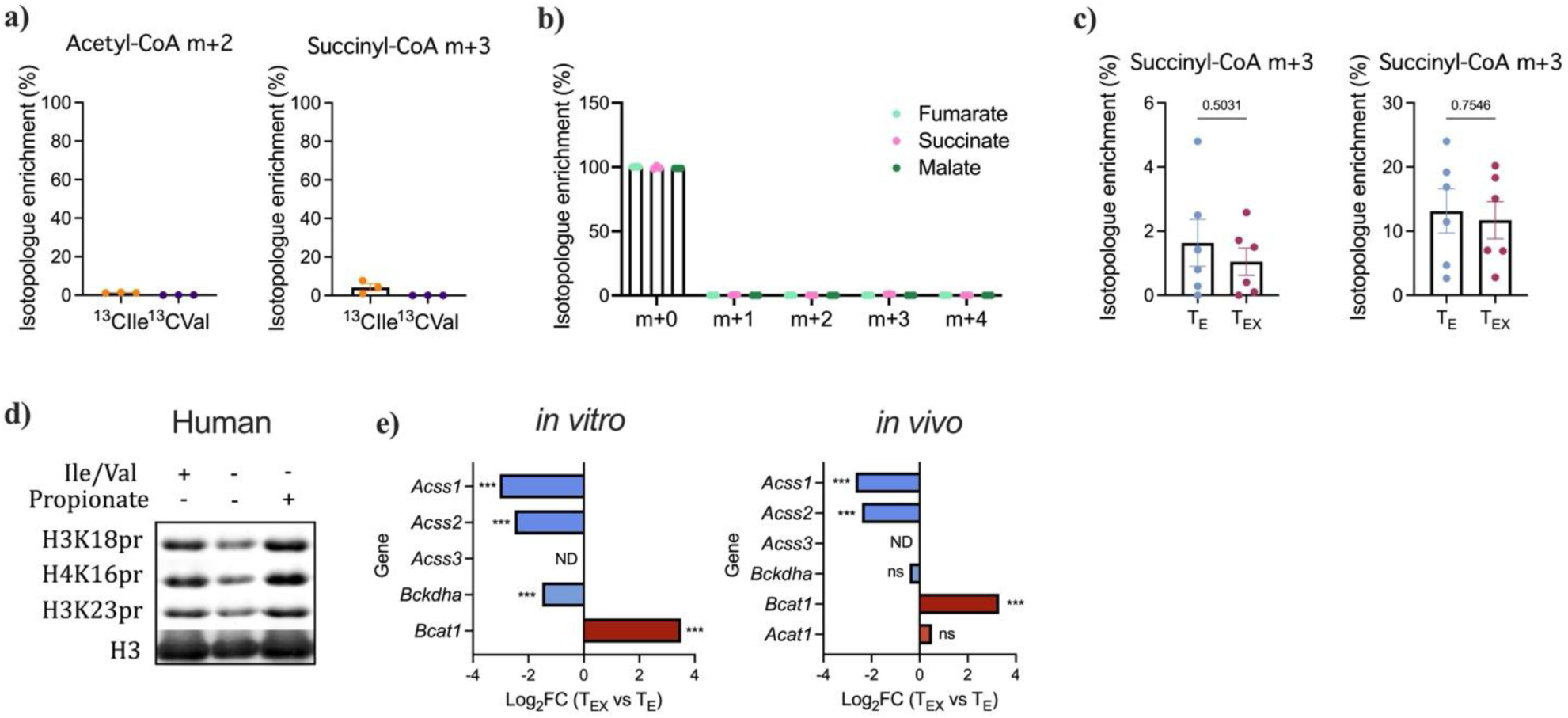
Histone Kpr is fueled by BCAA and propionate metabolism, which are altered during T cell exhaustion. **a)** Percent isotopic labeling of acetyl-CoA m+2 and succinyl-CoA m+3 following 2h tracing with [^13^C_6_]-isoleucine and [^13^C_5_]-valine in activated CD8^+^ T cells. **b)** Percent isotopic labeling (m+0 to m+4) of fumarate, succinate and malate following 2h tracing with [^13^C_6_]-isoleucine. **c)** Percent m+3 labeling of succinyl-CoA following 2h tracing with [^13^C_6_]-isoleucine and [^13^C_3_]-propionate in T_E_ and T_EX_ cells. **d)** Immunoblot of indicated histone marks upon Ile/Val deprivation and/or propionate supplementation in human CAR T cells. **e)** Log_2_ fold change of indicated acyl-CoA synthetase and BCAA catabolic genes in T_EX_ relative to T_E_ cells, derived from two independent RNA-seq datasets. Left: in vitro dataset comparing chronically versus acutely stimulated P14 CD8⁺ T cells at day 7 [40]. Right: in vivo dataset comparing PD-1^hi^ TIM-3^+^ TILs versus tumor dLN T cells from MC38-gp33 tumor-bearing mice [23]. *Acss3* was not detected (ND) in either dataset. Negative log₂FC indicates lower expression in T_EX_ cells. ***padj < 0.001; ns, not significant. Data are shown as mean ± SEM; statistical significance was determined by unpaired Student’s t-test (c).

**Fig. S3.**
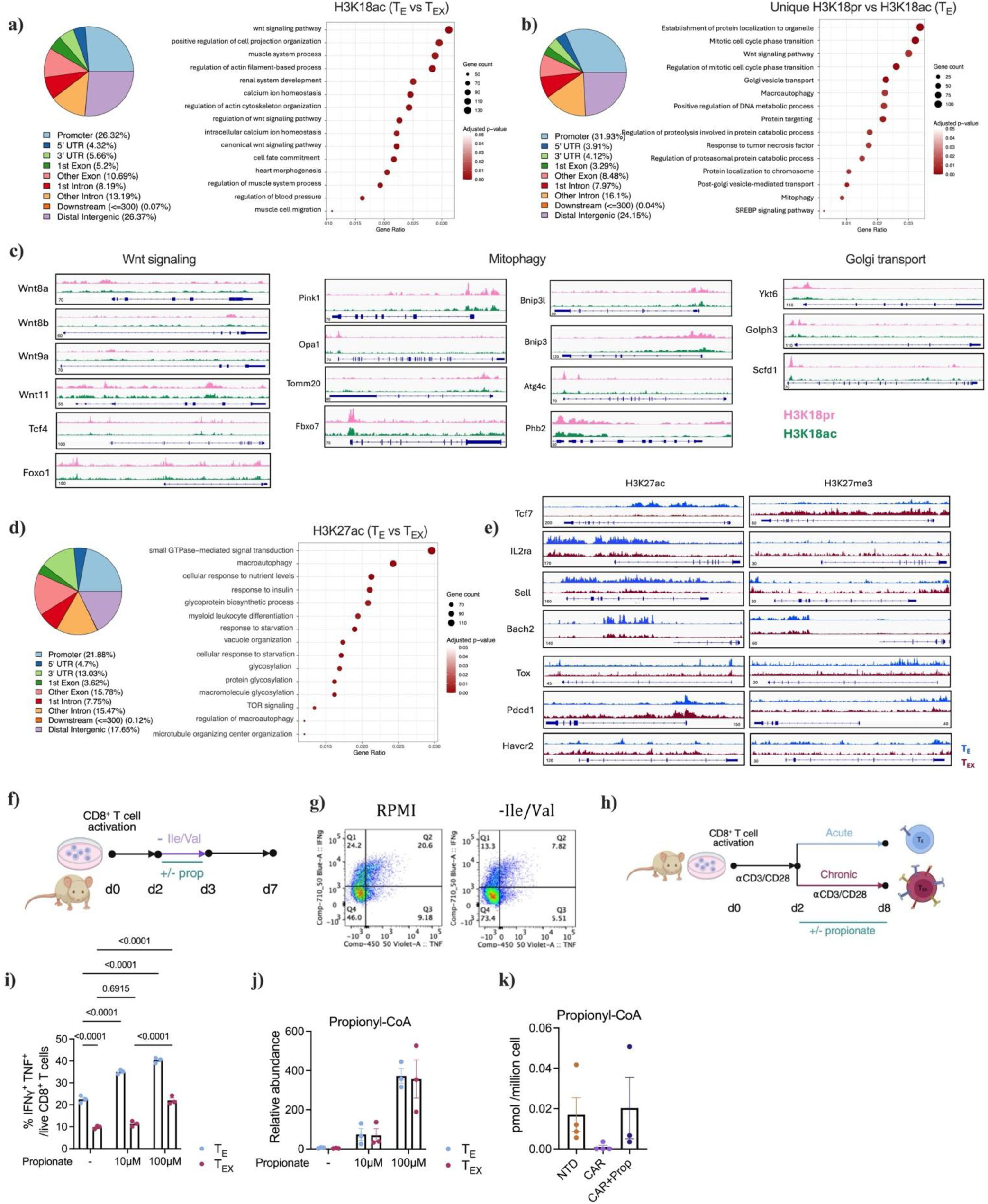
Nutrient conditions control histone propionylation and T cell fate and function. **a)** Genomic distribution of H3K18ac peaks in T_E_ cells and gene ontology analysis of H3K18ac peaks unique in T_E_ cells. **b)** Genomic distribution of H3K18pr peaks lacking overlap with H3K18ac in T_E_ cells, and gene ontology analysis of H3K18pr-unique peaks. **c)** Genome tracks of indicated loci showing H3K18pr and H3K18ac occupancy in T_E_ cells. **d)** Genomic distribution of H3K27ac peaks in T_E_ cells and gene ontology analysis of the H3K27ac peaks unique in T_E_ cells. **e)** Genome tracks of indicated loci showing H3K27ac and H3K27me3 in T_E_ and T_EX_ cells. **f)** CD8^+^ T cells were isolated and activated with αCD3/CD28 and IL-2. At day 2, cells were cultured under Ile/Val deprivation conditions with or without 100 µM propionate for 24 hours, before being returned to complete RPMI for an additional 4 days. **g)** Representative dot plot of IFNγ^+^ and TNF^+^ CD8^+^ T cells under fed (RPMI) or Ile/Val-deprived (-Ile/Val) conditions following restimulation. **h)** CD8^+^ T cells were isolated and activated with αCD3/CD28 and IL-2. Two days post activation, cells were either cultured without αCD3/CD28 to generate T_E_ cells, or restimulated with αCD3/CD28 every other day until day 8 to generate T_EX_ cells. Both T_E_ and T_EX_ cells were cultured with or without 10 or 100 µM propionate from day 2 to day 8. **i and j)** IFNγ^+^ and TNF^+^ production (i) and relative pr-CoA abundance (j) in T_E_ and T_EX_ cells following 6-day supplementation with 10 or 100µM propionate. **k)** Relative pr-CoA abundance in non-transduced (NTD) T_E_ and in HA-28z CAR T_EX_ cells cultured with or without propionate (100 µM) as described in Fig. 1j. Data are shown as mean ± SEM; statistical significance was determined by one-way (k) or two-way ANOVA with Tukey’s multiple-comparison test (i, j).

**Fig. S4.**
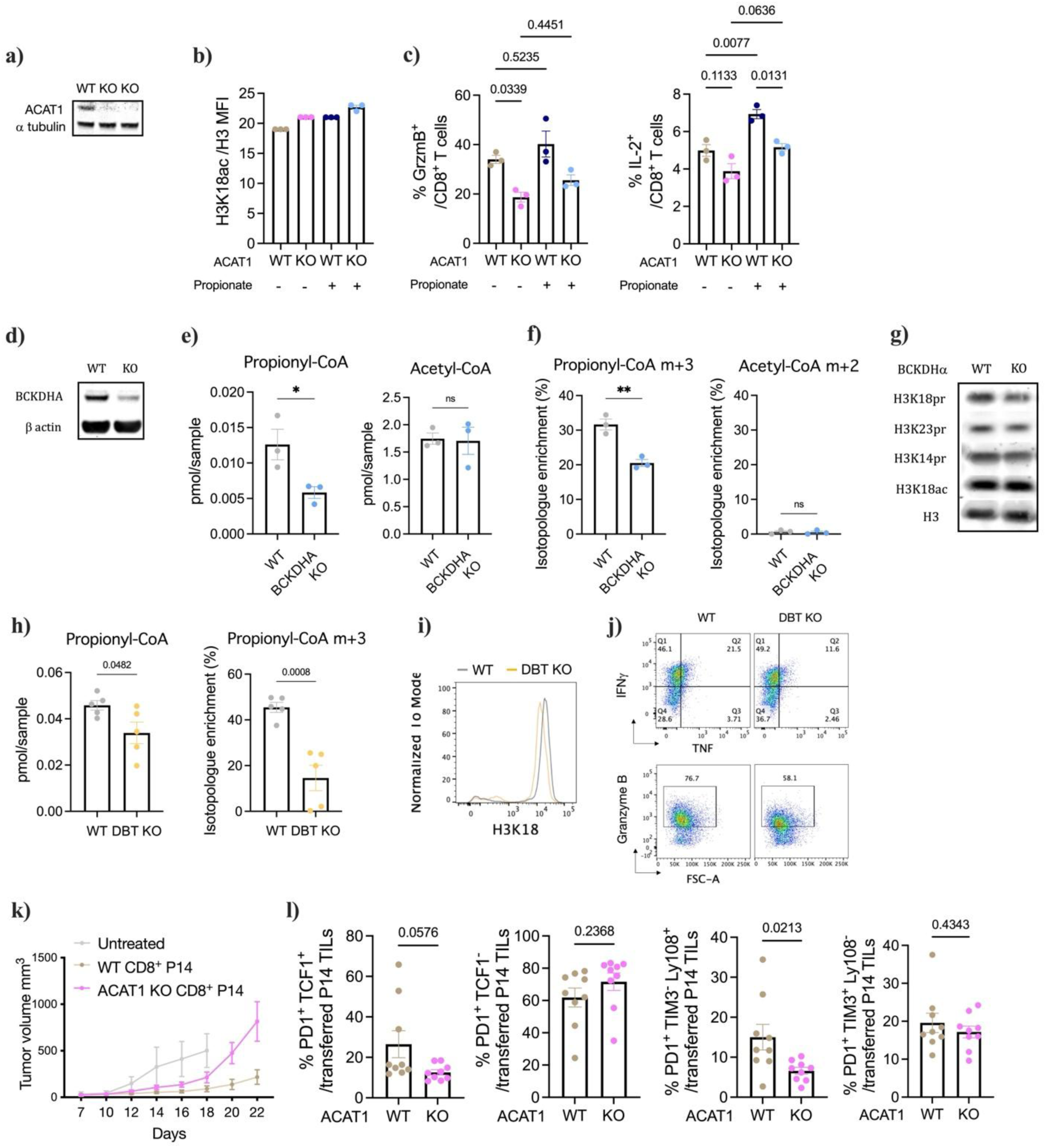
BCAA and propionate metabolism regulate T cell Kpr, fate and function. **a)** ACAT1 protein expression in WT and KO CD8^+^ T cells. **b)** H3K18ac levels normalized to total H3 in Rosa control (WT) and ACAT1-deficient (KO) P14 CD8^+^ TILs. **c)** Percentage of GzmB^+^ and IL-2^+^ cells among Rosa control (WT) and ACAT1-deficient (KO) CD8^+^ T cells, cultured with or without 100 µM propionate following restimulation. **d)** BCKDHA protein expression in WT and KO CD8^+^ T cells. **e)** pr-CoA and acetyl-CoA abundances (pmol per million cells) in BCKDHA WT and KO CD8^+^ T cells. **f)** Percent m+3 labeling of pr-CoA and m+2 labeling of acetyl-CoA following 2h tracing with [^13^C_6_]-isoleucine in BCKDHA WT and KO CD8^+^ T cells. **g)** Immunoblot of indicated histone marks in BCKDHA WT and KO CD8^+^ T cells. **h)** pr-CoA abundance (pmol per million cells) and percent labeling of pr-CoA m+3 in DBT WT and KO CD8^+^ T cells. **i)** Representative histograms of H3K18pr staining in DBT WT and KO CD8^+^ T cells, measured by flow cytometry. **j)** Percentage of IFNγ^+^ and TNF^+^ and GzmB^+^ CD8^+^ T cells among DBT WT and KO CD8^+^ T cells following restimulation. **k)** Tumor growth curve of mice bearing MC38-gp33 tumors and transferred with Rosa control (WT) or ACAT1-deficient (KO) P14 CD8^+^ T cells. **l)** Percentage of PD1^+^ TCF1^+^; PD1^+^ TCF1^-^; PD1^+^ TIM3^-^ Ly108^+^ ; PD1^+^ TIM3^+^ Ly108^-^ cells among Rosa control (WT) and ACAT1-deficient (KO) P14 TILs. Data are shown as mean ± SEM; statistical significance was determined by unpaired Student’s t test (e, f, h, l) and by one-way ANOVA with Tukey’s multiple-comparison test (c).

**Fig. S5.**
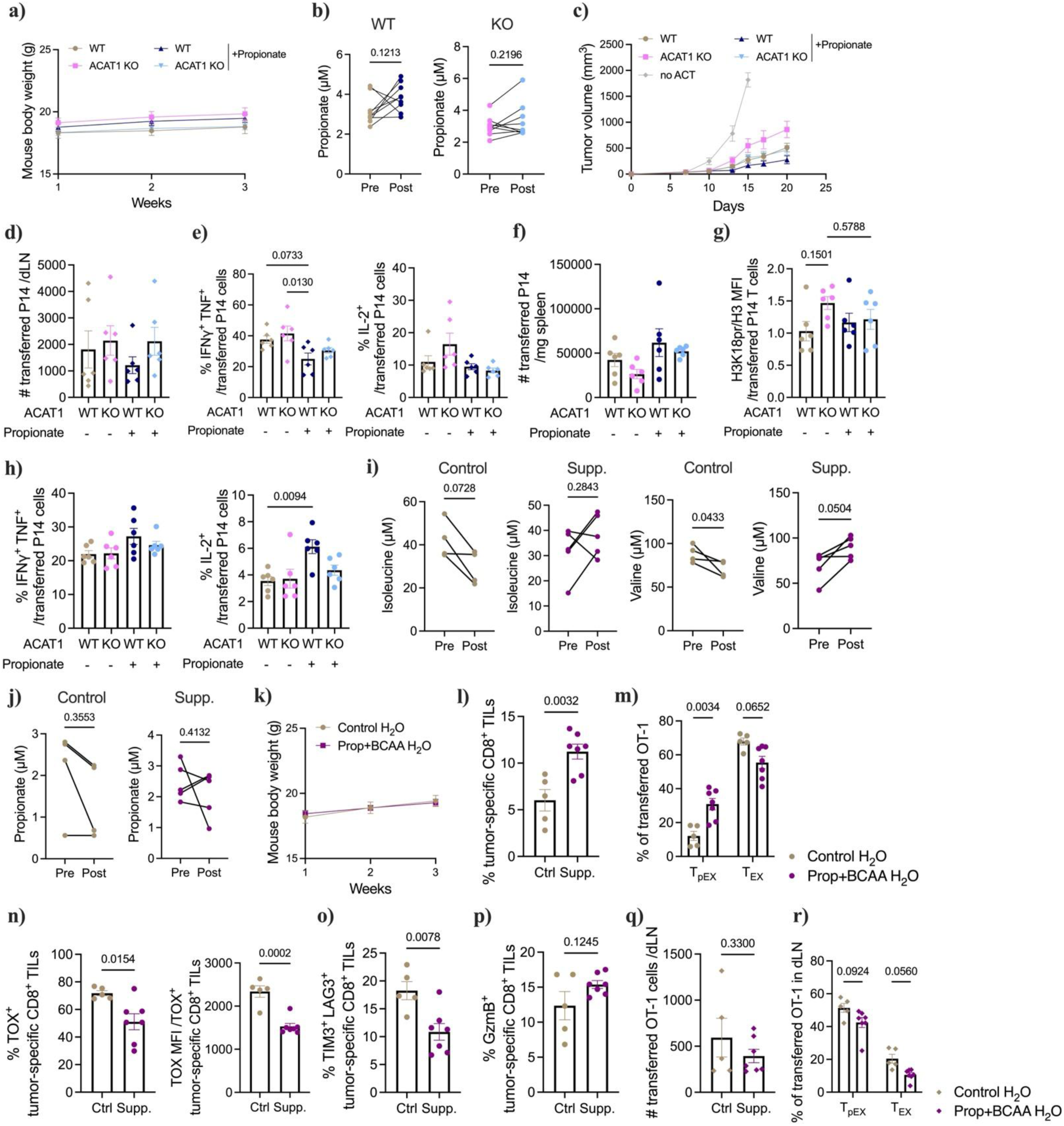
Nutrient composition in microenvironments dictates T cell fate and function via histone propionylation. **a)** Mouse body weight in indicated groups. **b)** Concentrations of propionate in plasma of MC38-gp33 tumor-bearing mice measured by LC-MS. **c)** Tumor growth curve of mice bearing MC38-gp33 tumors and transferred with Rosa control (WT) or ACAT1-deficient (KO) P14 CD8^+^ T cells, receiving control or propionate-supplemented drinking water. **d)** Numbers of P14 CD8^+^ T cells per tumor dLN in indicated groups. **e)** Percentage of IFNγ^+^ and TNF^+^ and IL-2^+^ cells among P14 CD8^+^ T cells from the tumor dLN of indicated groups following *ex vivo* restimulation. **f)** Numbers of P14 CD8^+^ T cells per mg of spleen in indicated groups. **g)** H3K18pr levels normalized to total H3 in P14 CD8^+^ T cells from spleen of indicated groups. **h)** Percentage of IFNγ^+^ and TNF^+^ and IL-2^+^ cells among P14 CD8^+^ T cells from spleen of indicated groups upon *ex vivo* restimulation. **i)** Concentrations of isoleucine and valine in plasma of MC38-gp33 tumor-bearing mice measured by LC-MS. **j)** Concentrations of propionate in plasma of MC38-gp33 tumor-bearing mice measured by LC-MS. **k)** Mouse body weight in indicated groups. **l)** Frequency of OT-1 CD8^+^ TILs in control and supplemented mice. **m)** Percentage of progenitor exhausted (PD1^+^ TCF1^+^ T_pEX_) and terminally exhausted (PD1^+^ TCF1^-^ T_EX_) OT-1 TILs in control and supplemented mice. **n)** Percentage and MFI of TOX^+^ cells among OT-1 TILs of control and supplemented mice. **o)** Percentage of TIM3^+^ LAG3^+^ cells among OT-1 TILs of control and supplemented mice. **p)** Percentage of GzmB^+^ cells among OT-1 TILs of control and supplemented mice. **q)** Numbers of OT-1 T cells per tumor dLN in control and supplemented mice. **r)** Frequency of T_pEX_ and T_EX_ OT-1 T cells in the tumor dLN of control and supplemented mice. Data are shown as mean ± SEM; statistical significance was determined by unpaired (l, n-q) or paired (b, i, j) Student’s t test, and by one-way (d-h) or two-way (m, r) ANOVA with Tukey’s multiple-comparison test.

